# Perinatal Exposure to Metal Mixtures Disrupts Neuronal Function and Behavior

**DOI:** 10.1101/2025.09.05.673216

**Authors:** Naveen Chandra, Benyamin Karimi, Ankita Bhobe, Susmitha Pagadala, Shekhar Pandey, Sylvia S. Sanchez, Bonnie Yeung-Luk, Emily Rivera, Mark J. Kohr, Beth Riess, Keeve E. Nachman, Shyam Biswal, Fenna C. M. Sillé, Austin R. Graves

## Abstract

**Background:** Environmental exposure to heavy metals such as lead (Pb), arsenic (As), hexavalent chromium [Cr(VI)], and cadmium (Cd), (PACC), is linked to neurodevelopmental disorders. These metals often co-occur in contaminated environments, but their combined effects on brain development remain poorly understood.

**Objective:** To test the hypothesis that perinatal exposure to a mixture of environmentally relevant levels of Pb, As, Cd, and Cr(VI), causes developmental defects in cognition, behavior, and neuronal function.

**Methods:** Female C57BL/6J mice were exposed to either a single metal or the PACC mixture in drinking water. Exposure began two weeks preconception and continued until weaning at postnatal day 21. Juvenile mice were tested at 4–5 weeks of age in open field (locomotion), novel object recognition (short-term memory), Y-maze (working memory), and elevated plus maze (anxiety-like behavior). A subset of animals underwent Whole-cell patch-clamp recordings in the medial prefrontal cortex (mPFC) and hippocampal CA1 neurons.

**Results:** Perinatal exposure to PACC metal mixture increased anxiety-like behavior and impaired short-term memory but not locomotion or working memory. Pyramidal neurons in mPFC and hippocampal CA1 displayed increased intrinsic excitability, mPFC neurons also showed elevated amplitude in spontaneous excitatory postsynaptic currents.

**Discussion:** Our findings suggest that perinatal exposure to the PACC metal mixture impairs cognition, increases anxiety-like behavior, and alters neuronal function in specific brain regions of juvenile mice, leading to disruption in neuronal function and behavior later in life. Further studies are needed to provide mechanistic insight into how perinatal heavy metal exposure affects neuronal development.

**Highlights:**

- Perinatal exposure to a metal mixture including lead (Pb), arsenic (As), hexavalent chromium (Cr(VI)), and cadmium (Cd), collectively termed PACC metal mixture—impairs cognition and increases anxiety in mice.
- Neuronal excitability and synaptic transmission are altered in medial prefrontal cortex after PACC metal mixture exposure.
- PACC mixture exposure decreases short-term memory in both males and females, and increases anxiety in males
- Principal component and clustering analyses reveal that PACC mixture exposure and control mice form distinct, nonoverlapping populations in physiological-behavioral space.
- Environmentally relevant PACC metal mixtures exert stronger effects than individual metals alone.

## Introduction

Heavy metal pollution remains a persistent and escalating worldwide environmental and public health concern. Because heavy metals and metalloids are elemental substances, they cannot be degraded by natural processes ^1^. They enter the environment through both natural events—such as soil erosion, forest fires, and volcanic activity, —as well as through anthropogenic activities including mining, smelting, manufacturing of batteries, pesticide application, and fossil fuel combustion ^2, 3^. Human exposure can occur via ingestion, inhalation, consumption of contaminated water, and dermal contact ^4^. In many settings, heavy metals occur as complex mixtures in the environment, —such as those frequently detected at U.S. Superfund sites ^5–7^—that may exert additive or synergistic toxic effects ^8–10^. Among environmental neurotoxicants, lead (Pb), arsenic (As), cadmium (Cd), and hexavalent chromium [Cr(VI)], collectively referred to as the PACC metal mixture—rank among the top hazardous substances on the Agency for Toxic Substances and Disease Registry (ATSDR)’s Priority List, occupying the 1^st^, 2^nd^, 7^th^, and 17^th^ positions, respectively ^11^.

Each metal has been independently associated with neurodevelopmental deficits, including oxidative stress, mitochondrial dysfunction, neuroinflammation, and disruption of neurotransmitter systems ^12–23^. Pb, for example, impairs synaptic plasticity and memory, especially in children ^24, 25^, while As and Cd contribute to redox imbalance and inflammatory signaling in the brain ^16–22^. Cr(VI), while less extensively investigated in the context of neurotoxicity, has been reported to cause acute oxidative injury to brain tissue and behavioral abnormalities in animal models ^23^. However, most toxicological studies have examined these metals in isolation, leaving a critical gap in our understanding of the cumulative, synergistic, or antagonistic effects of real-world metal mixtures. Investigating the combined impact of these environmentally relevant metals during early brain development is therefore essential to model human exposure more accurately and reveal potential mixture-specific mechanisms of neurotoxicity.

In humans, the most dynamic neurodevelopmental processes occur in the last trimester of pregnancy through the first 2–3 years of life. Numerous studies have demonstrated that PACC metals can cross the placental barrier and accumulate in fetal tissues, including the brain ^26^. Therefore, developing fetuses and infants are among the most vulnerable groups to heavy metal exposure ^27, 28^. During the critical neurodevelopmental window, the central nervous system (CNS) undergoes intricate and highly regulated processes, including cell proliferation, differentiation, migration, apoptosis, synaptogenesis, and synaptic pruning ^29–31^. These tightly orchestrated developmental events are particularly sensitive to disruptions from environmental toxicants ^32, 33^. Even low-level exposure during critical windows can result in long-lasting or irreversible changes in brain structure and function, increasing the risk of neurodevelopmental disorders such as autism spectrum disorder (ASD), attention-deficit/hyperactivity disorder (ADHD), learning disabilities, and emotional dysregulation ^34–36^.

In addition to their neurodevelopmental effects, heavy metals affect neuronal function in the adult brain by altering mitochondrial activity, reducing ATP production, increasing reactive oxygen species (ROS), and inducing neuronal death through apoptotic or necrotic mechanisms ^37^. For instance, Pb disrupts mitochondrial structure and triggers ER stress via PINK1-mediated MFN2 ubiquitination ^38^. Pb also antagonizes NMDAreceptor activation, disrupting calcium signaling and synaptic plasticity ^39, 40^. Arsenic impairs complexes II and IV, leading to neuron degeneration and motor issues ^41^. Cr(VI) induces apoptosis through ROS-related mitochondrial damage ^42^, while Cd reduces ATP and increases oxidative stress ^43^. These disruptions compromise energy and glutamate signaling, especially in the mPFC and CA1, which altogether may manifest as pathological states associated with lifelong deficits in cognition, behavior, and general health. Real-world exposures rarely involve single metals. Instead, toxicants are often found as mixtures, where they may act additively, synergistically, or antagonistically, altering toxicity and bioavailability ^39, 40^. To test a model that better resembles real-world conditions, we exposed perinatal C57BL/6J mice using drinking water at U.S. EPA maximum contaminant levels (MCLs) for As, Cd, and Cr(VI) and/or the historical U.S. EPA action level (AL) for Pb ^44^, beginning two weeks before conception and continuing through weaning (postnatal day 21). We then assessed cognition- and anxiety-related behavior and functional changes in the mPFC and hippocampal CA1 ^45^. These regions are essential for memory, executive function, and behavioral flexibility ^46, 47^, and become increasingly interconnected after birth ^48, 49^. Their coordination supports adaptive learning and emotional regulation ^50, 51^, and disruption in either area—via disease or toxicant exposure—can impair these functions ^52–54^.

Based on the evidence, we hypothesized that perinatal exposure to the PACC metal mixture leads to behavioral impairments and functional changes in CA1 and mPFC. By studying how exposure to PACC metal mixtures in early development affects neural function and behavior, this study aims to provide mechanistic insight into how environmental metal mixtures impact neurodevelopment and identify potential neuronal circuits underlying the observed behavioral phenotypes. Our findings contribute to a growing understanding of how environmental exposures to heavy metal mixtures influence neurodevelopment and highlight the need for further research to inform public health considerations.

## Methods

### Animal Care and Metals Exposure Models

Wild-type C57BL/6J mice (Jackson Laboratories, Bar Harbor, ME, U.S.A.) were used in this study. All animals were housed in the Johns Hopkins School of Public Health vivarium, which follows the Animal Welfare Act regulations and Public Health Service (PHS) Policy (PHS Animal Welfare Assurance #: D16-00173 (A3272-01)) and is accredited by the Association for the Assessment and Accreditation of Laboratory Animal Care (AAALAC International # 000503). All experiments were performed according to NIH guidelines for the care and use of laboratory animals and were approved by the Institutional Animal Care and Use Committee of Johns Hopkins University (Protocol # MO22H379), following the National Research Council’s Guide for the Care and Use of Laboratory Animals. In addition, we followed the ARRIVE guidelines 2.0 ^55^ for the entire study.

All mice were maintained under a reversed 12-hour light-dark cycle (lights off at 9 AM, on at 9 PM), at 20–22°C and 55% ± 5% humidity, with ad libitum access to food (AIN-93M: Research Diets, New Brunswick, NJ, U.S.A., CAT# D10012Mi) and water (Crystal Geyser, CG Roxane, NY, U.S.A.). Food and water were replaced every 3–4 days, and averaged individual intake was tracked throughout the experiment. Prior to breeding, male mice were housed individually with environmental enrichment (extra paper bedding and mouse igloo) and females were group-housed (5/cage). Dams were housed individually post-mating within environmentally enriched cages.

Female mice received either individual metals or the PACC metal mixture beginning two weeks prior to timed pregnancies and continuing through gestation and lactation. F1 offspring were weaned on postnatal day 21 and subsequently provided standard water. Metals were dissolved in water at concentrations consistent with U.S. EPA-defined environmentally relevant levels. For arsenic (10 parts per billion – 10 ppb), cadmium (5 ppb), and hexavalent chromium (100 ppb), these reflect the maximum contaminant levels (MCLs) ^56^. Lead was included at 15 ppb, the previous U.S. EPA action level (AL), as it lacks an MCL ^44^. Since October 2024, the U.S. EPA has suggested a new “trigger level” for lead of 10 ppb as part of updated regulations ^57^.

**Figure 1A** illustrates the overall experimental timeline including exposure, breeding, weaning, behavioral assessments, and terminal electrophysiological recordings. **Figure 1B** shows the experimental groups and the specific concentrations of each metal used for exposure.

**Figure 1.**
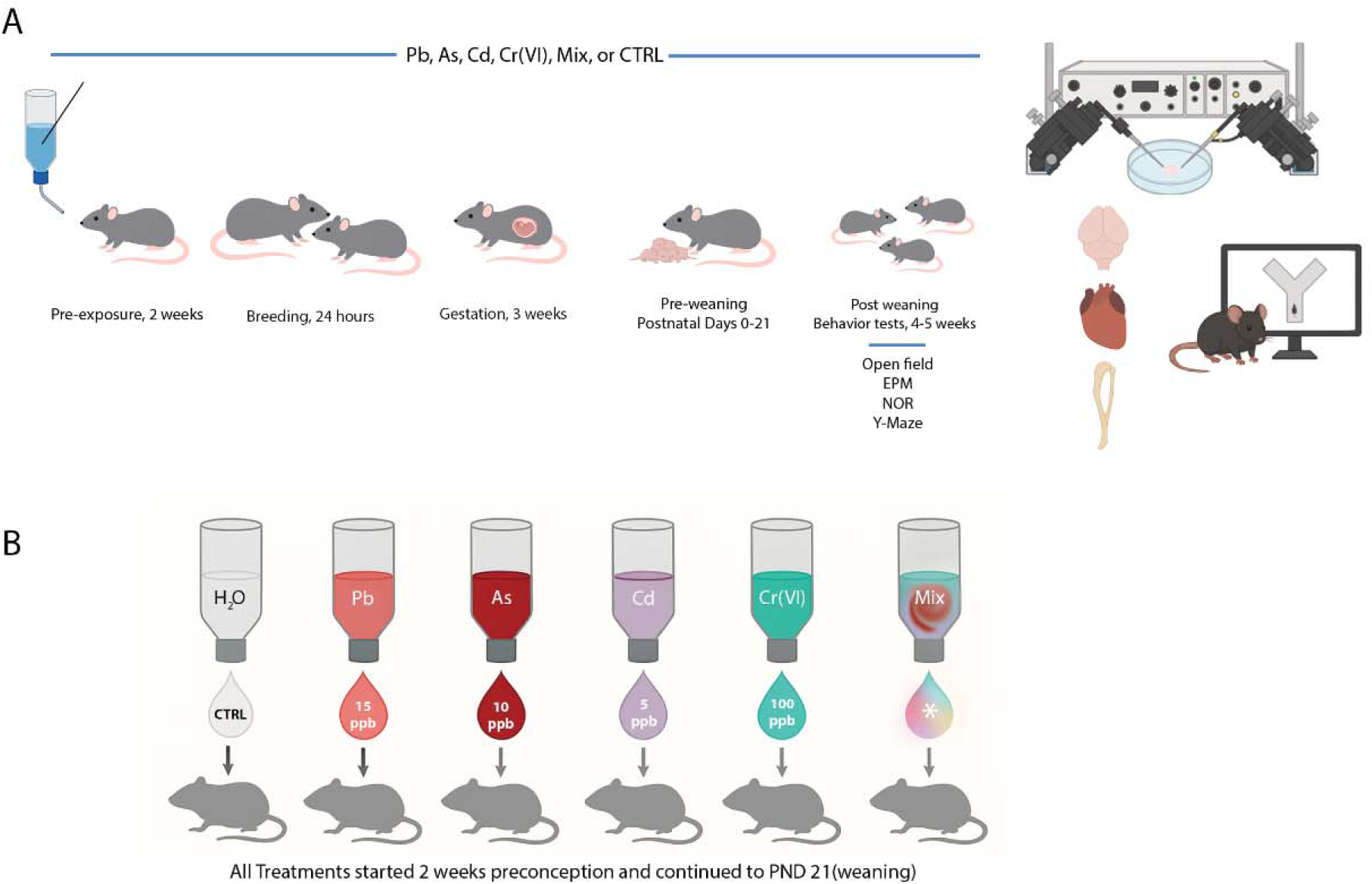
Experimental design for perinatal metal exposure and downstream assessments. **(A)** Timeline and experimental workflow. Female C57BL/6J mice were exposed to either individual metals (Pb, As, Cd, or Cr(VI)), a combined mixture (PACC; labeled as “Mix”), or control (CTRL) through drinking water starting 2 weeks before mating. Exposure continued throughout gestation (3 weeks) and lactation until postnatal day 21 (weaning). Offspring were tested at 4–5 weeks of age using a behavioral battery including Open Field (locomotor activity), Elevated Plus Maze (EPM; anxiety-like behavior), Novel Object Recognition (NOR; short-term memory), and Y-Maze (spatial working memory). After behavioral testing, tissues were collected for electrophysiology, cardiac assessment, and histological or molecular analysis. **(B)** Exposure groups included unexposed controls (CTRL), individual metal exposures (Pb, As, Cd, Cr), and the PACC metal mixture group (Mix). All metals were administered at maximum contaminant levels (MCLs) for As, Cd, and Cr(VI) ^56^, or the historical U.S. EPA action level (AL – 15 ppb, as it lacks an MCL) for Pb^#^ via drinking water ^44^. Offspring from each group were evaluated in the same behavioral and physiological pipeline. All behavioral and electrophysiological experiments were performed blind to exposure conditions. ^#^NOTE: Since October 2024, the U.S. EPA has suggested a new “trigger level” for lead of 10 ppb as part of updated regulations ^57^.

F1 juvenile mice were subjected to behavioral tests at 4-5 weeks of age to assess locomotion (Open Field Test), short-term memory (Novel Object Recognition), spatial working memory (Y-maze spontaneous alternation), and anxiety (Elevated Plus Maze) (n≥7 combined from 4 repeat experiments, ≥4males and ≥3 females/group). Behavior studies were followed by electrophysiological recordings from acute brain slices in a subset of the mice (unexposed controls and PACC metal mixture groups, n=8, 4 males and 4 females/group). In addition, hearts, liver and lung were excised and weighed, and tibia length assessed. Heart size was determined by normalizing heart weight-to-tibia length, which accounted for any age-dependent differences and^58^ and was used to rule out cardiac hypertrophy or atrophy as a confounding variable that could potentially affect behavioral or neurophysiological changes.

#### Sample Size

We used a total of 120 breeding pairs (20 pairs per group, 6 groups in total: Control, Pb, As, Cd, Cr(VI), and PACC metal mixture) across 4 waves of experiments. Out of the 20 breeding pairs in each group, 5 pairs of controls, 9 of mixtures, 2 of Pb, 3 of As, 8 of Cd, and 5 of Cr(VI) resulted in pregnancy. The resulting F1 offspring were used for behavioral and electrophysiological experiments.

#### Inclusion & Exclusion Criteria

We aimed to include at least 7 male and 7 female mice per group for behavioral testing. Mice were excluded from analysis if they showed signs of illness or injury prior to or during the study. For electrophysiological experiments, at least 8 animals (4 males, 4 females) from the control and PACC metal mixture groups were included. Cells with over 20% change in series resistance during recording were excluded from final analysis.

#### Randomization and Blinding

Dams were randomly assigned to exposure groups using a random number generator. F1 offspring were randomly assigned to behavioral testing days and order. All experiments were conducted blind to exposure group and analyses were automated and conducted blind to exposure group.

### Outcome Measures

Maternal weight, food and water intake were monitored throughout the study. Behavioral tests for anxiety (EPM), memory (NOR, Y-maze), and locomotion (OFT) were performed on all six groups. Electrophysiological analyses were limited to unexposed and PACC metal mixture groups.

### Experimental procedures

For a detailed description of the experimental procedures, refer to the **Supplementary Materials Section 1** (Supplemental Methods: Detailed Experimental Procedures).

### Behavioral Tests

#### Open Field Test (OFT)

The Open Field Test was performed to assess locomotor activity and exploratory behavior. Mice were placed in a 40 x 40 x 40 cm open arena for 5 minutes. Position and velocity were tracked via overhead IR-camera (Hotpet Optics, Shenzhen Ailipu Technology Co., Guangdong, China) using ANY-maze software (Stoelting, IL, U.S.A.). Each test was conducted under dim red light during the dark cycle, as illustrated in **Supplementary Fig. S2**.

#### Novel Object Recognition (NOR)

The purpose of the NOR test was to assess short-term recognition memory in mice. Mice explored a chamber (40 x 40 x 40 cm) containing two identical objects for 5 minutes (training phase), after which they were returned to the home cage. After 30 minutes, one object was replaced with a novel item and mice were returned for a 5-minute test phase. Exploration time of each object was recorded, and novel object preference was assessed by dividing the amount of time exploring the novel object, divided by the total time spent exploring the novel and familiar objects combined. Exploration of each object was defined as being within 5cm of the object, with its head oriented towards the object within an arc of 45 degrees in front of the mouse. Mice were tracked via overhead IR-camera (Hotpet Optics, Shenzhen Ailipu Technology Co., Guangdong, China) using ANY-maze software (Stoelting, IL, U.S.A.).

#### Y-Maze Spontaneous Alternation

The Y-maze spontaneous alternation test is a readout for working memory, drawing on the rodent preference for exploring new environments. The equipment consisted of three arms (80 cm long) 120° apart. Mice were placed in one arm and given 5 minutes free exploration time in the maze, under dim red light. Observation centered on whether the mice returned to the most recently entered arm, or alternated to return to a different, new arm. Mice were tracked via overhead IR-camera (Hotpet Optics, Shenzhen Ailipu Technology Co., Guangdong, China) using ANY-maze software (Stoelting, IL, U.S.A.). Quantification of spontaneous alternation was defined as previously described ^59^.

#### Elevated Plus Maze (EPM)

EPM is a widely used behavioral paradigm to assess exploratory drive and anxiety-like behavior in rodents. The apparatus featured two open arms (30 x 5 x 15 cm) and two enclosed arms (30 x 5 x 15 cm), elevated 50 cm above the ground, connected by a central platform (5 x 5 cm). Mice were placed in the center facing an open arm and allowed to explore for 5 minutes in dim red light. Time spent and entries into each arm were tracked using an overhead IR-camera (Hotpet Optics, Shenzhen Ailipu Technology Co., Guangdong, China) and quantified using ANY-maze (Stoelting, IL, U.S.A.).

### Electrophysiology

Electrophysiological recordings from hippocampal CA1 and mPFC pyramidal neurons assessed the effects of perinatal PACC metals exposure on intrinsic excitability and synaptic function. Parameters measured included resting membrane potential, input resistance, spike count in response to step current injections (−50 to +200 pA, 25 pA increments, 100 ms duration), and spontaneous EPSCs at −70 mV (analyzing amplitude, frequency, rise and decay kinetics). Whole-cell patch-clamp recordings used an Axon Instruments 700b amplifier and digitizer with pClamp acquisition (Molecular Devices, San Jose, CA, U.S.A.); data were analyzed in Clampfit 11.2 (Molecular Devices, San Jose, CA, U.S.A.). Recordings with >20% series resistance change were excluded. Solution compositions are provided in **Table S1**.

Within 3 days of behavioral experiments, mice were deeply anesthetized with inhaled isoflurane. Coronal slices of the brain (300 µm) were sectioned with a VT1200S Vibratome (Leica Microsystems, Buffalo Grove, IL, U.S.A.) in ice-cold oxygenated (95% O_2_/5% CO_2_) sucrose-based cutting solution. Slicing was started just posterior to the olfactory bulbs. Medial PFC, located in anterior slices, was recognized by its midline location and laminar organization in the medial cortex, before the emergence of anterior commissure. Dorsal CA1 was targeted in more posterior sections by its distinctive C-shaped pyramidal layer surrounding the dentate gyrus. Slices were incubated for 45 minutes at 32°C in standard artificial cerebrospinal fluid (ACSF, for details of the electrophysiological solutions see **Table S2)** and then held at room temperature in the oxygenated ACSF until recording.

Slices were visualized using a Nikon microscope with IR-DIC optics and a 40x water-immersion objective (Nikon Instruments, INC., Melville, NY, U.S.A.). Borosilicate pipettes (Sutter P1000, Sutter Instrument CO., Novato CA, U.S.A.) with an open-tip resistance of 4–6 MΩ were filled with a standard potassium-gluconate-based internal solution. Slices were perfused at 1–2 mL/min with heated (32°C) oxygenated ACSF. Both mPFC and hippocampus recordings targeted visually identified pyramidal neurons. Our recordings were focused on excitatory synaptic input due to the experimental design and technical considerations. Spontaneous EPSCs (sEPSCs) were recorded in normal ACSF with 100μm picrotoxin to block GABAergic transmission. We used a K-gluconate-based internal solution and at –70 mV holding potential, optimized for the estimation of glutamatergic transmission as well as intrinsic excitability within the same cell.

### Statistical Analysis

Analyses were performed in GraphPad Prism 10 (Graphpad Software, Boston, MA, U.S.A.) using mixed-effects models with the Geisser–Greenhouse correction for repeated-measures/partially paired designs; familywise error was controlled with multiplicity-adjusted post-hoc tests (Tukey or Šídák, as applicable), computing individual variances for each comparison (two-tailed α=0.05). To evaluate the effects of PACC metal mixture exposure on brain and behavior, a multivariate analysis was conducted using data from animals subjected to both behavioral and electrophysiological assessments (n=18; 10 PACC metal mixture, 8 control) ^60^. Input variables consisted of behavioral metrics from the Novel Object Recognition (NOR) and Elevated Plus Maze (EPM), as well as electrophysiological parameters from mPFC neurons: resting membrane potential, input resistance, intrinsic excitability (spike count in response to step current injection), and spontaneous excitatory postsynaptic current (sEPSC) amplitude, frequency, and decay kinetics. All variables were z-scored for normalization.

Principal Component Analysis (PCA) was utilized to reduce data dimensionality and identify covarying features across behavioral and electrophysiological dimensions within individual mice. The first two principal components explained 64% of the total variance. K-means clustering (k=2) was applied to the PCA-transformed dataset to determine natural groupings within the sample. Statistical significance of cluster separation was evaluated by calculating the summed Euclidean distance of each animal from its group centroid and comparing this to all 43,758 possible 10-vs-8 groupings.

To understand potential interactions in the PACC metal mixture, albeit with a single dose of each metal, we performed an abbreviated additivity analysis (**Supplemental Information: Section 3**). Briefly, we adopted a four-step approach for evaluating dose additivity in mixtures proposed by Hertzberg et al ^61^. We also applied a simple method for assessing response additivity using effect summation ^62, 63^.

To identify outliers, we used Grubbs’ test (GraphPad Prism 10, α = 0.05). Grubbs’ test was applied once per group to detect a single extreme value relative to the group mean and standard deviation.

## Results

### Maternal and Offspring Health Metrics Were Largely Unaffected by Heavy Metal Exposure

To establish a baseline for interpretation of the outcomes and ruling out possible confounding factors, we first determined if dam or offspring health were impacted by exposure with individual metals or PACC metal mixture. Maternal weight gain and consumption of food and water during the gestation period were not significantly different between control and treatment groups. Cannibalization, although observed, was rare and not dependent on exposure (**Supplementary Fig. S1**).

To investigate the systemic effects of metal exposure beyond the brain, we examined cardiac development in offspring using the normalized heart weight-to-tibia length (a measure of body size) (**Supplementary Fig. S4**). We did not observe any significant variations in heart size in the offspring exposed to either the PACC metal mixture or individual metals compared to the controls. Although cadmium-treated males suffered slight decreases in heart weight-to-tibia length ratio, but this seemed to be secondary to a reduction in total body weight. The findings indicate that exposure to environmentally relevant doses of perinatal metals does not induce a gross physiological toxicity that would confound subsequent behavioral or functional analyses.

### Perinatal Metal Exposure Does Not Impair Locomotor Activity

As intact gross motor skills are crucial for performing all the behavioral tests in this study, we performed an open field test (OFT) to determine if any of the observed behavioral changes were due to nonspecific shifts in activity level or locomotion. Neither single metal nor PACC metal mixture exposure affected the total distance traveled, compared to control untreated animals. This finding supports the idea that general locomotion and arousal were not affected by PACC metals treatment. Model assumptions were validated using QQ plots of residuals (**Supplementary Fig. S3**), confirming the normal distribution of all measured behavioral and electrophysiological properties, and thus the appropriateness of mixed-effects modeling.

### PACC Metal Mixture Exposure Impaired Recognition Memory but Not Spatial Working Memory

Heavy metals are known neurotoxicants that can interfere with brain development. Exposure to heavy metals, especially during prenatal period ^64^, can negatively affect neurotransmission ^9, 65, 66^, increase oxidative stress ^16–23^, and change neuronal excitability ^56^, which can impair memory and learning and lead to increased anxiety ^67, 68^.

To determine the effect of perinatal metal exposure to PACC metal mixture and the individual metals on short-term memory, we used the Novel Object Recognition (NOR) test. NOR is used to assess recognition memory in rodents and takes advantage of the rodents’ spontaneous exploratory preference for novel objects.

In male mice, PACC metal mixture exposure significantly reduced novel object preference, compared to controls (**Fig. 2A**, CTRL vs. Mix, *p*=0.0099). Same trend was observed in females (**Fig. 2B**, *p*<0.0001). Moreover, a significant reduction in novel object preference was also observed in female As group (*p*=0.0476) and Cr (*p*=0.0311). Pooling sexes showed an even more pronounced reduction in novel object preference in PACC metal mixture exposure compared to control (*p*<0.0001). In addition, reduced novel object preference was observed in the pooled As (*p*=0.0264) and Cd (*p*=0.007) versus the pooled control (**Fig. 2C**). These observations indicate that perinatal exposure to the PACC metal mixture impaired short-term recognition memory.

**Figure 2.**
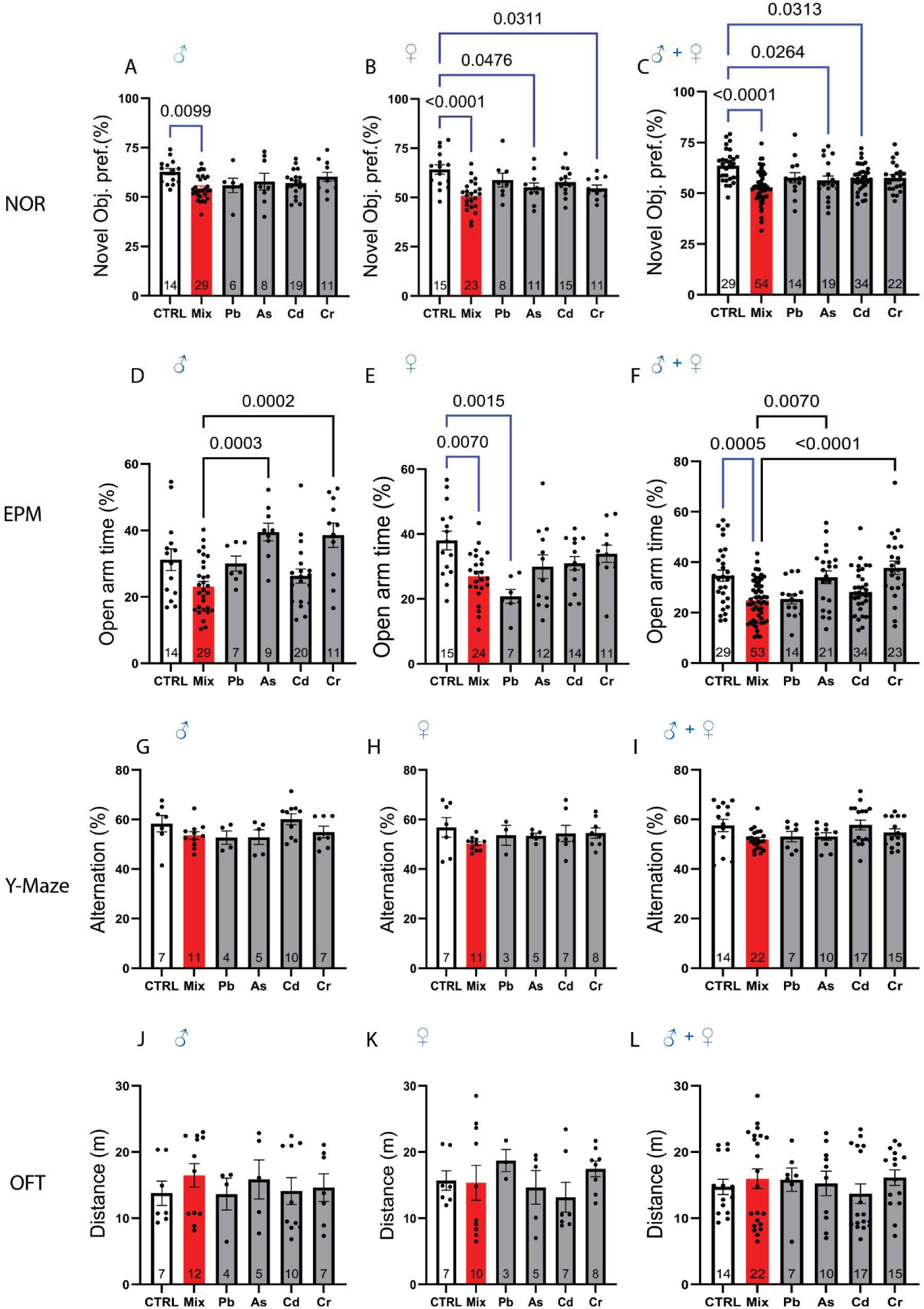
Perinatal exposure to PACC metal mixture alters behavioral performance. (**A–C**) Novel Object Recognition (NOR) test assessing short-term recognition memory in (**A**) male, (**B**) female, and (**C**) all mice combined. PACC metal mixture-exposed mice (labeled “MIX”) showed significantly reduced novel object preference compared to untreated controls (“CTRL”), indicating impaired short-term memory. In some comparisons, individual metals such as Pb and Cd also reduced performance, but PACC metal mixture had a more consistent and stronger effect across sex. (**D–F)** Elevated Plus Maze (EPM) test measuring anxiety-like behavior in (**D**) male, (**E**) female, and (**F**) all mice combined. PACC metal mixture exposure significantly decreased open arm time, indicating heightened anxiety. While some individual metals such as Pb also reduced open arm time, the effect of PACC metal mixture was generally more pronounced. Open-arm exploration time in PACC metal mixture-treated mice was also significantly lower than individual metals, again supporting the notion of combined toxicity exceeding that of individual metals. **(G–I)** Y-Maze spontaneous alternation, assessing spatial working memory, in (**G**) male, (**H**) female, and (**I**) all mice combined. No significant differences were observed across groups, indicating that PACC metal mixture and individual metals did not impair spatial working memory under the tested conditions. **(J–L)** Open Field Test measuring general locomotor activity in (**J**) male, (**K**) female, and (**L**) all mice. Total distance traveled did not differ significantly between control, PACC metal mixture, or any of the individual metal groups, suggesting that the observed behavioral deficits in NOR and EPM were not due to changes in gross motor function or arousal. Data are presented as mean ± SEM, with each dot representing one mouse. One-way ANOVA with multiple comparisons was used for statistical analysis. Blue significance bars denote comparisons between the untreated control group and each exposure group (PACC metal mixture or individual metals). Black significance bars denote comparisons between the PACC metal mixture group and each individual metal group. Exact *p*-values are shown above each line. The numbers at the bottom of each bar graph denote the number of animals per group (n≥7 combined from 4 repeat experiments, ≥4males and ≥3 females/group).

### Anxiety-Like Behavior is Increased in Offspring Exposed to the PACC Metal Mixture

To examine the impact of perinatal metal exposure on anxiety-like behavior, we employed the Elevated Plus Maze (EPM), a widely used test that relies on rodents’ natural aversion to open and elevated spaces. The time spent by male mice in the open arm showed a reduced albeit unsignificant trend, while PACC metal mixture-exposed female mice showed a similar significant trend (*p*=0.007). In addition to the PACC metal mixture group, lead-exposed females also spent significantly less time in the open arms than controls (*p*=0.0015) (**Fig. 2E**). When sexes were pooled, PACC metal mixture exposed mice displayed, reduced open arm time relative to control (*p*=0.0005), as did As exposure (*p*=0.007) (**Fig. 2F**). These data indicate that perinatal exposure to the PACC metal mixture or to individual metals such as As can promote anxiety-like behavior. **Supplemental Tables S3 and S4** present detailed behavioral outcomes, including sex-stratified NOR and EPM data, plus group OFT and Y-maze comparisons.

### Perinatal PACC Metal Mixture Exposure Enhanced Neuronal Excitability in mPFC

Emotional regulation, memory, and learning require uninterrupted systematic communication between cortical and subcortical regions, most notably, mPFC and hippocampus. Both of these regions have been found to be vulnerable to insults during neurodevelopment. To explore how perinatal PACC metal mixture exposure altered intrinsic and synaptic neuronal functions in these areas, we performed whole-cell patch-clamp recordings from neurons in mPFC and CA1 in acute slices of mice following behavioral testing. We measured resting membrane potential (RMP) as an indicator of baseline excitability, input resistance as an indicator of responsiveness of the membrane to inputs, and number of spikes (action potentials) in response to increasing steps of injected currents as a measure of intrinsic neuronal excitability.

In dorsal CA1 pyramidal neurons, neither resting membrane potential (RMP) nor input resistance differed significantly in neurons of PACC metal mixture-exposed mice compared to that of the controls (**Fig. 3A–B**). Although PACC metal mixture-exposed CA1 neurons had a tendency toward greater spike output to current steps, this did not reach significance at any step (**Fig. 3C**). In the mPFC, PACC metal mixture-exposed neurons showed depolarized RMP (*p*=0.0174; **Fig. 3F**), while input resistance was not altered. PACC metal mixture-exposed mPFC neurons did not discharge more action potentials at several steps of current compared to controls (**Fig. 3G and H**).

**Figure 3.**
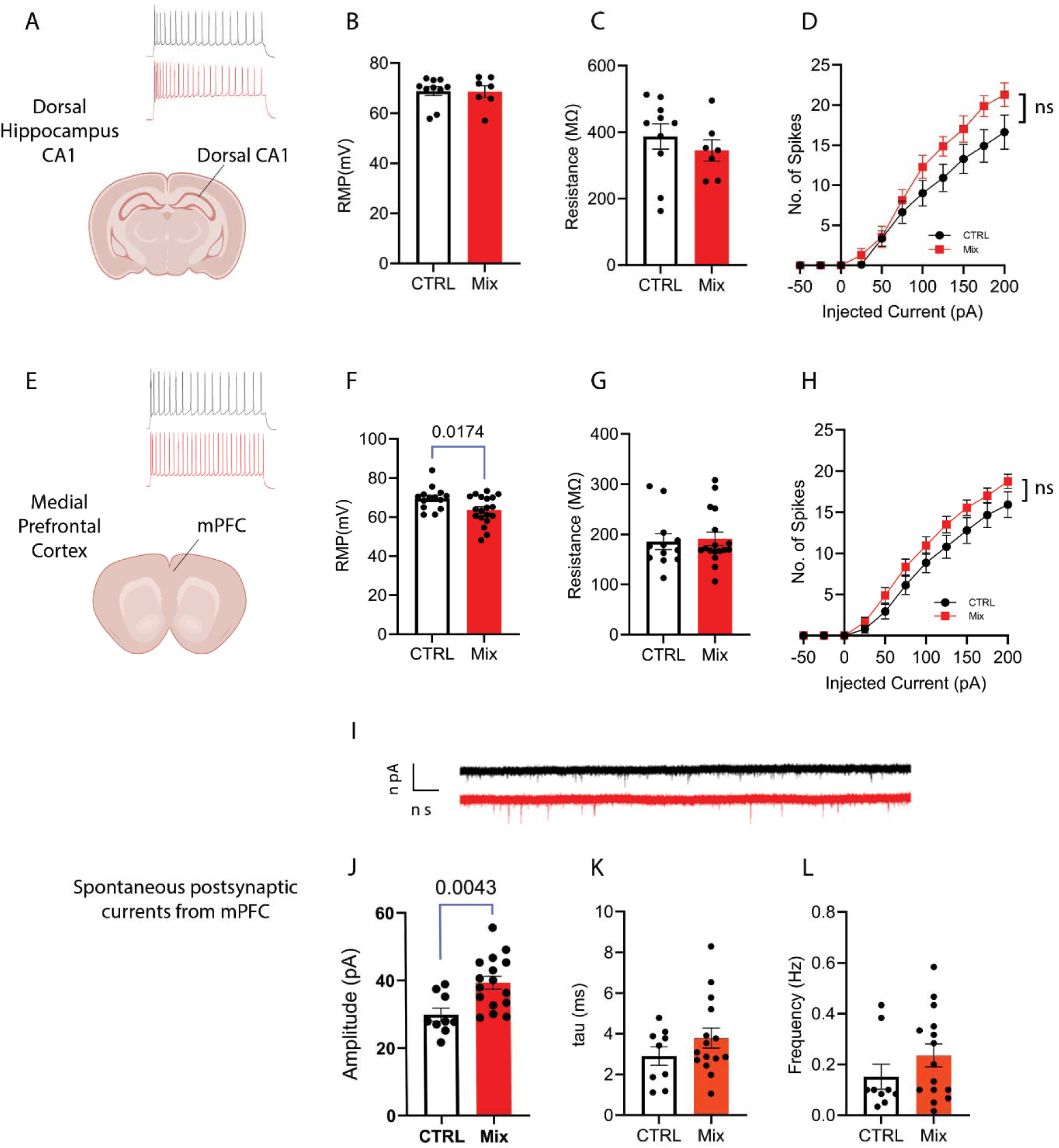
Perinatal exposure to PACC metal mixture alters intrinsic excitability and synaptic transmission in hippocampus dCA1 and mPFC. (**A-D**) Dorsal Hippocampus CA1, (**A**) Representative traces for dorsal CA1. (**B**) Resting membrane potential (RMP) did not differ between control (“CTRL”) and PACC metal mixture-exposed groups (“Mix”). (**C**) Input resistance (Rin) was also unchanged. (**D**) Spike counts during stepwise depolarizing current injection (–50 to +200 pA) showed a non-significant trend toward increased excitability in the PACC group. Top left: The traces show representative action potential firing patterns recorded from CA1 pyramidal neurons in CTRL (black) and PACC metal mixture-exposed (red) mice during depolarizing current steps (**E**) Representative traces for Medial Prefrontal Cortex (mPFC), (**F**) PACC metal mixture exposure caused a significant depolarization of RMP (*p*=0.0174), indicating increased baseline excitability. (**G**) Input resistance remained unchanged between groups. (**H**) Spike output in mPFC was not significantly increased in PACC metal mixture-exposed neurons at multiple current steps, although a non-significant trend toward increased excitability in the PACC metal mixture group was observed. (**I**) Representative traces for Spontaneous Excitatory Postsynaptic Currents (sEPSCs) in mPFC. (**J**) PACC metal mixture exposure significantly increased sEPSC amplitude (*p*=0.0043), suggesting enhanced postsynaptic response to excitatory inputs. (**K**) No significant difference in decay kinetics (tau). (**L**) Frequency of sEPSC events was not significantly altered, indicating that presynaptic release probability or event rate was unaffected. Total number of mice: 8, 4 males and 4 females/group. Data are expressed as mean ± SEM. Statistical significance determined using unpaired t-tests for single-point comparisons and two-way repeated-measures ANOVA with Sidak’s multiple comparisons test for spike-frequency analysis.

Despite no change in intrinsic spike output, mPFC showed a depolarized resting membrane potential, suggesting altered synaptic drive. Therefore, we recorded spontaneous postsynaptic currents (sEPSC) from mPFC pyramidal neurons to quantify the characteristics of excitatory synaptic input. To this end, we assessed amplitude as an indication of the magnitude of individual synaptic events, since larger amplitudes reflect stronger postsynaptic depolarization per input. SEPSC frequency was used to estimate how often spontaneous presynaptic release occurred. Finally, sEPSC decay time constant (tau) was used to measure the duration of synaptic events, which can influence how long each synaptic event lasts and influences the temporal summation of excitatory synaptic inputs. PACC metal mixture exposure significantly increased sEPSC amplitude (*p=*0.0043; **Fig. 3G**), indicating enhanced postsynaptic excitability, likely due to increased expression or function of postsynaptic conductances. Frequency and tau were not significantly different between groups (**Fig. 3H–I**). These results indicate that perinatal PACC metal mixture exposure selectively enhanced postsynaptic responsiveness to excitatory input in mPFC neurons, without altering presynaptic release frequency or synaptic kinetics.

### PCA and Clustering Reveal Distinct Behavioral-Electrophysiological Signatures of PACC Metal Mixture Exposure

To integrate behavioral and physiological findings across multiple domains, we performed Principal Component Analysis (PCA) using six features: NOR performance (time investigating novel object, divided by total time spent investigating novel and familiar objects combined), EPM behavior (time in open arm), resting membrane potential, input resistance, neuronal excitability (number of spikes evoked by +200pA current injection), and sEPSC amplitude. The first two principal components (PC1 and PC2) together accounted for 64% of the total variance in the dataset (**Fig. 4A**). When plotted in two-dimensional PCA space, mice segregated clearly by exposure status, with PACC metal mixture-exposed animals forming a distinct cluster from controls (**Fig. 4B**).

**Figure 4.**
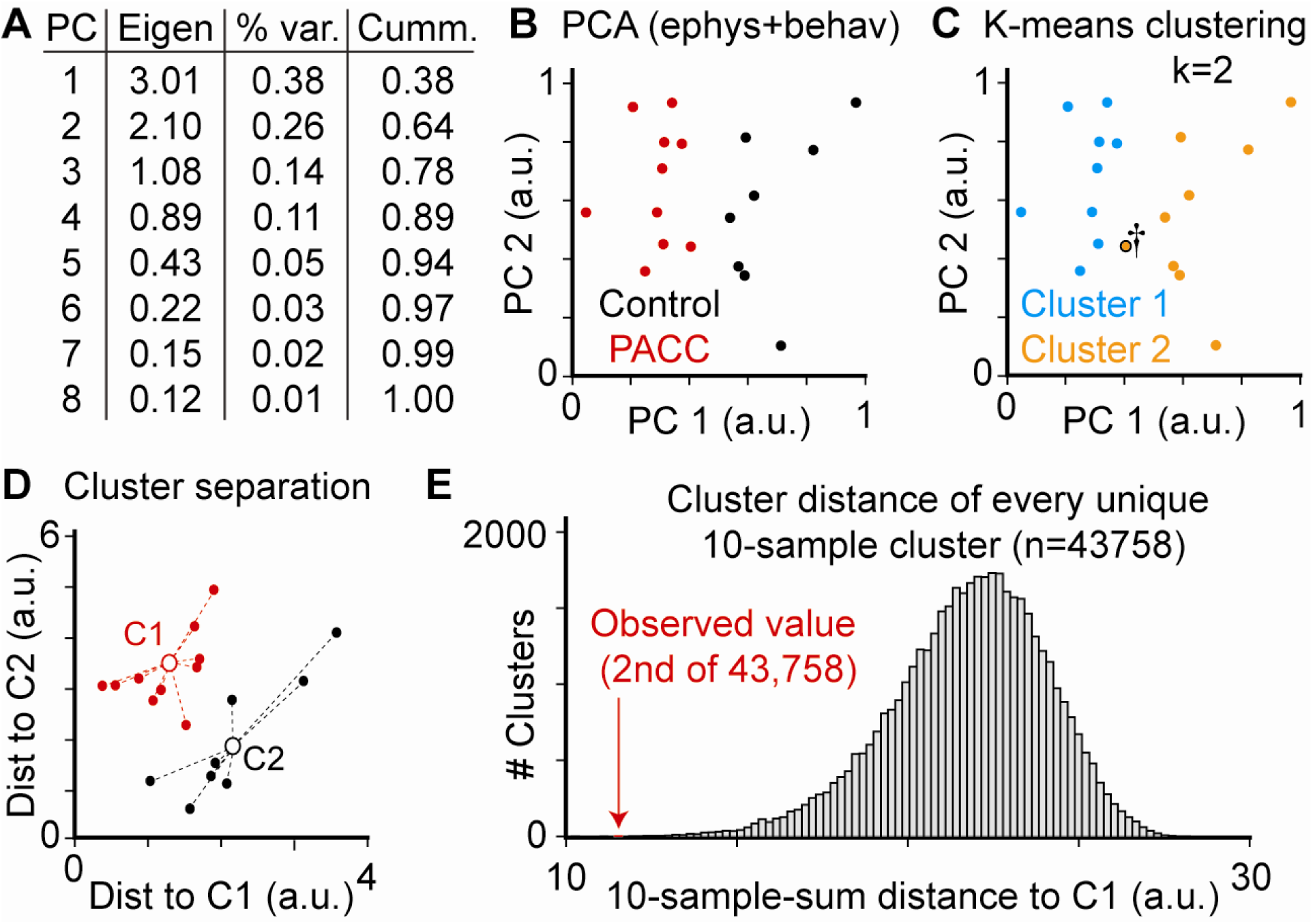
of behavioral and electrophysiological data reveals significant differences between PACC Metal Mixture and untreated mice. **(A)** Table of each Principal Component (PC), its Eigenvalue, variance explained by each individual PC, and the cumulative variance explained by including each successive PC. **(B)** Plot of all data (n=18 cells) for first two PCs, colored by exposure group (red is PACC metal mixture, n=10; black is untreated control, n=8). **(C)** Unsupervised K-means clustering into 2 groups (k=2) divides experimentally observed data into two very similar clusters as separated by exposure condition, with only a single cell misassigned (marked by dagger). **(D)** Plotting each cell as a function of distance from its cluster center (either C1 or C2, respectively) revealed significant separation between clusters (*p*<0.05 1-way ANOVA). **(E)** Histogram shows distribution of all possible distances to all possible cluster centers. Experimentally observed clustering (red) is the second lowest summed distance (i.e., second tightest clustering) of all possible permutations, supporting the extreme unlikelihood that observed clustering occurred by chance.

Unsupervised K-means clustering (k=2) recapitulated this division with remarkable accuracy, misclassifying only a single neuron into the other treatment condition (**Fig. 4C**). To assess the statistical robustness of this separation, we performed permutation analysis of all 43,758 possible 10-to-8 cell clusterings (the sample sizes from each experimental condition). The experimentally observed clustering exhibited the second smallest total Euclidean distance to its centroid among all permutations, underscoring the improbability of such separation arising by chance (**Fig. 4D–E**).

These multivariate analyses corroborate the conclusion that perinatal PACC metal mixture exposure induces a coherent and distinct neurobehavioral phenotype marked by increased anxiety, impaired memory, and altered neuronal function.

### Perinatal Metal Exposure Does Not Alter Normalized Heart Size in Offspring

To examine a potential link between neurocognitive impairment and heart disease, as has been reported ^69^ we next examined heart size as a potential measure of pathological heart growth in male and female offspring exposed to the PACC metal mixture or individual metals. We found that the heart weight-to-tibia length ratio—a widely accepted manner to normalize heart weight to total body size ^58, 70, 71^—did not change upon exposure to the PACC metal mixture compared to non-exposed male or female offspring, or with individual exposure to lead, arsenic, cadmium or chromium. However, hearts from female offspring exposed to chromium were significantly larger than hearts from female offspring exposed to the PACC metal mixture, but again neither group was significantly different from non-exposed female offspring. Overall, we found no evidence that PACC metal mixture exposure affected heart size (**Supplementary Fig. S4)**.

## Discussion

Perinatal exposure to environmental toxicants poses a critical risk to neurodevelopment. While previous studies have examined the impact of individual metals such as lead (Pb), arsenic (As), cadmium (Cd), or chromium [Cr(VI)] on brain function ^12–25^, relatively few have evaluated the effects of metal mixtures ^72–77^, despite their co-occurrence in the environment, particularly at highly contaminated U.S. Superfund sites ^5–7^. A substantial body of research has identified associations between exposure to heavy metal mixtures and neurocognitive as well as behavioral outcomes in humans and animals ^72–77^. Epidemiological studies indicate that children and adolescents exposed to these metals, particularly those residing near industrial areas, have a higher risk of reduced intelligence quotient (IQ), developmental delays, and psychomotor deficits ^78, 79^. Animal studies show that both single and combined exposures can result in cognitive impairments, neurobehavioral changes, and alterations in neurotransmitter systems. Some research describes synergistic neurotoxic effects and structural brain changes, especially involving dopaminergic and serotonergic pathways ^69, 80–82^.

However, there are gaps in the literature: many studies use artificial or single-metal exposures rather than actual mixtures, often do not focus on early developmental periods, and generally separate behavioral from neurophysiological evaluations while lacking detailed electrophysiological analysis of brain regions such as the hippocampus and prefrontal cortex. To address these limitations, this study examines perinatal exposure to a specified mixture of Pb, As, Cd, and Cr(VI) at environmentally relevant concentrations, focusing on periods of gestation and lactation. The study integrates behavioral tests with electrophysiological measurements to link observed cognitive and emotional effects to specific functional changes in the hippocampus and prefrontal cortex. Results indicate that exposure to realistic metal mixtures can lead to persistent changes in neuronal excitability and synaptic function, with implications for neurodevelopment. These findings provide mechanistic evidence regarding the cumulative effects of metal mixtures during critical developmental stages, highlighting the importance of evaluating combined exposures to inform public health guidelines and neurotoxicological understanding.

### Effects of Individual Metals and PACC Metal Mixture on Behavior

Consistent with previous work ^82–86^, we found that individual metal exposures exert modest effects on behavior. However, PACC metal mixture exposure induced more pronounced behavior impairments, namely reduced short-term memory, assessed using the NOR test and increased anxiety-like behavior on the elevated plus maze. Notably, locomotion (assessed by the open field test) and spatial working memory (assessed by the Y-maze test) were not impaired by the PACC metal mixture, indicating that its exposure selectively impacts neural processes implicated in affective and mnemonic processing, but not general arousal or motor coordination.

The fact that individual metals exerted subtler effects—while the full PACC metal mixture elicited robust behavioral deficits—raises the possibility of additivity, synergistic or antagonistic toxicity. For instance, both Pb ^87–89^ and As ^90–93^ can disrupt synaptic function independently, but their co-exposure might lead to compounded oxidative stress, impaired neurotransmission, or amplified inflammation in key brain regions ^80, 94^. This aligns with observation that individual metals produced only modest or fluctuating behavioral effects, whereas the full PACC metal mixture produced robust and reproducible recognition memory and anxiety-like impairments, suggesting that combined exposure leads to emergent toxicity not observed with exposures to individual metals. Importantly, the behavioral alterations we observed in PACC metal mixture exposed mice were further accompanied by dramatic alterations in neuronal function, specifically, increased excitability and larger sEPSC amplitudes in mPFC neurons, observed in the PACC metal mixture-exposed group alone. Furthermore, our multidimensional analyses (**Fig. 4)** support a correlation between the observed deficits in behavioral performance and electrophysiological function, suggesting that these deficits are linked, and that both deficits are likely downstream from the same pathogenic pathways initiated by exposure to PACC metal mixture. While electrophysiological data were not derived from individual metals, the lack of meaningful behavioral impairments in these groups suggests that synaptic and intrinsic changes observed in the PAC metal mixture group must be, or at least contribute in large part to, the basis for the cognitive and affective phenotypes observed. These data raise the possibility that perinatal PACC metal mixture exposure activates additive or perhaps synergistic processes to interfere with function of neurons within circuits relevant to memory and anxiety regulation.

### Electrophysiological Correlates of Behavioral Impairments

To clarify the functional underpinning of the observed behavioral impairment, we carried out whole-cell patch-clamp recordings in the mPFC and hippocampus dorsal CA1 regions (**Fig. 3**), two brain areas essential for cognitive function and emotional regulation ^95, 96^. Mouse neurons exposed to PACC metal mixture had substantially elevated intrinsic excitability in the mPFC. In mPFC, we also observed that neonatal exposure to PACC metal mixture was associated with a depolarized resting membrane potential and significantly increased sEPSC amplitude, which would further contribute to hyperexcitability via intrinsic and postsynaptic mechanisms, respectively. In mPFC, we observed no change in sEPSC frequency, suggesting that neonatal exposure to PACC metal mixture probably exerts its effect through a postsynaptic mechanism

These electrophysiological alterations closely parallel the behavior impairments in PACC metal mixture-exposed mice, i.e. enhanced anxiety and disruption of short-term, episodic memory. The mPFC plays a key role in regulating memory and affective behavior due to its top-down modulation of hippocampal and limbic systems ^97, 98^.

Although effective operation of mPFC is needed for normal executive function and behavioral inhibition, its hyperactivation, particularly by pyramidal cells is pathologic ^99^. Hyperexcitability of mPFC networks creates temporally incoherent and overactive activity patterns that compromise the neural coding necessary for novelty recognition and emotional regulation ^100–102^. Proper coordination of neural activity between the hippocampus and mPFC is crucial for episodic memory, and hyperexcitability can impair memory performance by reducing network synchrony and interfering with encoding or retrieval of novel stimuli ^100, 103^. Dorsal hippocampus-mPFC pathway is central for fear memory as tested by inhibitory avoidance task ^104^. Therefore, in anxiety-like situations, overactivation of mPFC efferents may lead to exaggerated anxiety responses, possibly through insufficient top-down modulation of subcortical regions such as the hippocampus and the amygdala.

Our findings are consistent with evidence that enhanced mPFC excitability during development caused by early-life stress, inflammation, genetic disruption, or toxicant exposure can precipitate behavioral dysfunction later in life ^105, 106^. Rodent models have demonstrated that hyperactivity of mPFC pyramidal neurons leads to both anxiety-like phenotype and memory impairments ^107, 108^, but interventions that restore mPFC function by decreasing its activity can reverse these phenotypes ^109^. Human imaging research also indicates that abnormal mPFC patterns of activity throughout childhood and adolescence correlate with greater anxiety and compromised emotion regulation ^110, 111^, which implies that the developmental course of mPFC function needs to be properly controlled to generate typical outcomes.

At the synaptic level, the observed increased sEPSC amplitude in PACC metal mixture-exposed mice most likely indicates increased postsynaptic expression of AMPA receptors, as well as posttranslational modifications that would increase AMPA receptor function. Such alterations are commonly observed in developmental stress or excitotoxic exposure models and are both associated with adaptive and maladaptive prefrontal circuit remodeling ^112, 113^. Whether this elevated excitatory drive in our model represents a secondary response to a prior insult or an early maladaptive process leading to pathology is still to be established.

### Additive, Synergistic, or Antagonistic Effects?

An important consideration in interpreting our results is whether the PACC metal mixture effects are indicative of additive, synergistic, or antagonistic interactions. Although our study was not specifically designed to differentiate between these possibilities, the fact that PACC metal mixture exposure led to more severe behavioral impairments, most notably in memory and anxiety, than exposure to any individual metal suggests that the effects of the PACC metal mixture reflect some level of interaction. To assess interaction, dose-response analyses with multiple doses of the metals are required. However, to understand potential interactions in the PACC metal mixture, albeit with a single dose of each metal, our abbreviated additivity analysis suggested less than additive mixture effects or antagonism (**Supplemental Information: Section 3, including Tables S5-S8**). When future studies with multiple doses allow for modeling of dose-response curves for the PACC metals, additivity analyses can be repeated and compared to this simple assessment. Since each metal can generate ROS or impair calcium homeostasis individually ^114, 115^, combined exposure may interfere with normal developmental signaling pathways and jointly lead to shared neurotoxic endpoints such as altered synaptic plasticity, glial activation, and impaired mitochondrial function.

### Cardiac Assessment and Neurodevelopment

Exposure to PACC metal mixture or heavy metals in general, can interfere with cardiac physiology which in turn may influence neurodevelopment ^116^. Some metals in the PACC metal mixture have been associated with cardiotoxicity through oxidative stress, vascular dysregulation, and interference with ion homeostasis ^116–118^. Abnormal cardiac development or function could alter cerebral perfusion or metabolic support during stages of brain maturation ^119, 120^. In our cohort, perinatal PACC metal mixture exposure did not measurably alter cardiac size in a way that could account for the observed neuronal and behavioral phenotypes; more sensitive functional assays (e.g., echocardiography, ECG, or serum biomarkers) would be required to exclude subtle effects.

### Mechanistic Insights, Future Directions, and Relevance to Human Health

Our research suggests a mechanistic understanding of how exposure to environmentally relevant mixtures of heavy metals during the perinatal period could disrupt neurodevelopment. Enhanced neuronal excitability and increased sEPSC amplitude in the mPFC suggest ion channel dysfunction, glutamatergic transmission, or compositional abnormalities of synaptic proteins. The PCA analysis revealed grouping of behavioral and electrophysiological phenotypes by exposure condition, indicating an integrated neurotoxic signature for PACC metal mixture exposure.

Follow-up experiments will expand on these results by carrying out transcriptomic profiling and other molecular analysis of the mPFC and hippocampus to discern changes in gene expression and downstream molecular pathways that are correlated with ion channel control, oxidative stress, and synaptic plasticity.

Since metal concentrations employed are representative of current U.S. EPA “safe” drinking water standards, our results are directly applicable to actual exposure settings. Children could be exposed to comparable PACC metal mixtures at levels permitted under current U.S. EPA drinking water standards during critical windows of brain development. Phenotypes observed here, cognitive deficit, heightened anxiety, and changed neuronal excitability, are similar to the phenotypes observed in neurodevelopmental disorders like attention deficit hyperactivity disorder (ADHD) and autism spectrum disorder (ASD) ^121, 122^. These results highlight the necessity of accounting for cumulative exposures and mixture toxicity in public health policy, rather than focusing solely on single pollutants. To better protect susceptible populations, EPA should incorporate evidence from metal mixture toxicity studies when revising drinking water regulations. The recent application of a hazard index to mixtures of per- and polyfluoroalkyl substances in drinking water illustrates a possible approach ^123^. A similar strategy could be considered for PACC metal mixtures should evidence of a common mode of action or neurological effects continue to accumulate.

## Acknowledgements

We acknowledge Thomas Hartung as an MPI on the grant supporting this research.

## Declaration of Generative AI and AI-assisted Technologies in the writing process

During the preparation of this work, the authors used AI-assisted technologies, namely PubMed’s Best Match and Query expansion features for information seeking; Grammarly to check grammar and spelling; and ChatGPT4o to generate pictographs for the schematic figures. After using these tools, the authors substantially reviewed and edited the content to ensure accuracy, relevance, and adherence to academic standards. The authors affirm that they take full responsibility for the content of the final manuscript.

## Conflict of interest statement

The authors claim no conflict of interest other than receiving financial support through a grant by the National Institute of Neurological Disorders and Stroke (NINDS), grant number RF1NS130672 (MPIs: F.C.M.S., S.B., K.E.N., T.H., Co-I: A.S.).

## Funding Statement

This research was supported by the National Institute of Neurological Disorders and Stroke (NINDS) grant number RF1NS130672 (MPIs: F.C.M.S., S.B., K.E.N., T.H., Co-I: A.S.). The content is solely the responsibility of the authors and does not necessarily represent the official views of the National Institutes of Health.

## CRediT authorship contribution statement

**Naveen Chandra:** Investigation; Data curation; Formal analysis; Visualization; Writing – original draft. **Benyamin Karimi:** Data curation; Formal analysis; Visualization; Writing – original draft; Writing – review & editing. **Ankita Bhobe:** Investigation. **Susmitha Pagadala:** Investigation. **Shekhar Pandey:** Investigation; Formal analysis. **Sylvia S. Sanchez:** Investigation. **Bonnie Yeung-Luk:** Investigation. **Emily Rivera:** Investigation. **Mark Kohr:** Investigation; Formal analysis. **Beth Riess:** Formal analysis. **Keeve E. Nachman**: Conceptualization; Funding acquisition; Supervision. **Shyam Biswal:** Conceptualization; Funding acquisition; Supervision. **Fenna C. M. Sillé:** Conceptualization; Investigation; Resources; Funding acquisition; Supervision; Project administration; Writing – original draft; Writing – review & editing. **Austin R. Graves:** Investigation; Formal analysis; Resources; Supervision; Project administration; Writing – original draft; Writing – review & editing.

## Notes

### Competing Interest Statement

The authors have declared no competing interest.

