## Supplemental Information for "Perinatal Exposure to Metal Mixtures Disrupts Neuronal Function and Behavior"

#### Supplementary Information

This file contains 3 sections: **1)** Supplemental Methods: Detailed Experimental Procedures, **2)** Supplemental Figures (S1-4) and Supplemental Tables (S1-8), and **3)** Method for Abbreviated Additivity Analysis for "*Perinatal Exposure to Metal Mixtures Disrupts Neuronal Function and Behavior*".

### 1. Supplemental Methods: Detailed Experimental Procedures

#### 1.1. Animal Husbandry and Experimental Design

All procedures adhered to the *ARRIVE 2.0 guidelines* [1]. Wild-type C57BL/6J mice (Jackson Laboratories, Bar Harbor, ME) were used. Mice were housed in the Johns Hopkins School of Public Health AAALAC-accredited animal facility (Accreditation #000503), with IACUC approval under protocol #MO22H379 and PHS Animal Welfare Assurance D16-00173 (A3272-01).

Mice were maintained on a reversed 12-hour light/dark cycle (lights off at 9:00 AM and on at 9:00 PM) at a stable temperature of 20–22°C and humidity of 55% ± 5%. Cages were individually ventilated with HEPA filtration and contained corncob bedding, a Nestlet square, and an igloo for enrichment. Mice were provided ad libitum access to water and AIN-93M diet (Research Diets, CAT# D10012Mi), with fresh food and water replaced every 3–4 days. During each change, individual food and water intake were measured for each dam using pre-weighed bottles and pellets, and intake was calculated based on weight change corrected for evaporation.

Male breeders were singly housed at least 1 week prior to mating, and female mice were group-housed 5 per cage until placed with males. Breeding pairs were housed together for up to 5 days. Plugged females were then single-housed and maintained under identical environmental conditions throughout gestation and lactation.

#### 1.2. Exposure Protocol

The PACC metal mixture consisted of lead (Pb), arsenic (As), cadmium (Cd), and hexavalent chromium (Cr(VI)) at EPA maximum contaminant levels (Pb: 15 ppb, As: 10 ppb, Cd: 5 ppb, Cr(VI): 100 ppb). Compounds were prepared using reagent-grade salts in ultrapure water and stored in light-protected bottles at 4°C. Final concentrations were verified by ICP-MS.

Exposures began two weeks prior to mating and continued through gestation and lactation. Control mice received Crystal Geyser bottled water without added metals. Treated dams were randomized to receive either a single metal or the PACC metal mixture. All F1 pups were weaned at postnatal day (PND) 21 and switched to untreated water. **Table S1**, summarizes Animal husbandry and metal exposure conditions

**Table S1. Animal husbandry and metal exposure conditions**

| **Category** | **Component** | **Details** |
|---|---|---|
| **Mouse Strain** | Strain | Wild-type C57BL/6J |
| | Vendor | Jackson Laboratories |
| **Housing Conditions** | Temperature | 20–22 °C |
| | Humidity | 55% ± 5% |
| | Light/Dark Cycle | Reversed 12 h light / 12 h dark (lights off at 9:00 AM) |
| | Enrichment | Paper bedding and igloos |
| **Food and Water** | Diet | AIN-93M (Research Diets, CAT# D10012Mi), ad libitum |
| | Water | Crystal Geyser (CG Roxane), changed every 3–4 days |
| | Intake Monitoring | Individual food and water intake measured during pregnancy/lactation |
| **Metal Exposure** | Exposure Duration | 2 weeks preconception through weaning (PND 21) |
| | Delivery Method | Via drinking water |
| | Groups | Control, Pb, As, Cd, Cr(VI), PACC metal mixture |
| | Pb Concentration | 15 ppb (action level) |
| | As Concentration | 10 ppb (maximum contaminant level) |
| | Cd Concentration | 5 ppb (maximum contaminant level) |
| | Cr(VI) Concentration | 100 ppb (maximum contaminant level) |
| **Breeding Design** | Breeding Pairs per Group | 20 pairs |
| | Number of Successful pregnancies | Wave totals per group<br>– Control: 5<br>– PACC metal mixture: 9<br>– Pb: 2<br>– As: 3<br>– Cd: 8<br>– Cr: 5 |

##### 1.3. Behavioral Testing Procedures

Behavioral tests were conducted from 10 AM to 3 PM under dim red light during the mice's dark cycle. Mice were habituated for one hour before each session, and apparatuses were cleaned with 70% ethanol between subjects. Data were collected using ANY-maze software (Stoelting Co.) and analyzed blind to treatment. **Figure S1** illustrates the behavioral tests assessing locomotion, anxiety-like behavior, and cognitive function.

###### 1.3.1. Open Field Test (OFT)

General locomotor activity and anxiety-related behavior in juvenile mice following perinatal exposure to individual heavy metals (Pb, As, Cd, Cr) or their mixture (PACC) were assessed using the Open Field Test (OFT). The protocol was adapted from established procedures [2,3] with modifications to accommodate the age and behavioral characteristics of the mice in this study.

Behavioral assessments were conducted during the dark phase of a reversed light cycle (9:00 AM–9:00 PM lights off), under dim red illumination (625–670 nm wavelength) to reduce stress and reflect natural activity patterns. Mice were acclimated in the testing room for at least one hour prior to assessment to limit transport-related stress effects.

The apparatus consisted of a square, opaque acrylic arena (40 × 40 cm, 40 cm walls) situated within a sound-attenuating enclosure to reduce external noise and visual interference. Overhead video tracking was accomplished with a near-infrared camera (Stoelting ANY-maze system, Wood Dale, IL), and the arena's center and periphery regions were digitally designated for analysis.

Each mouse was placed in the arena's center and permitted to explore freely for five minutes, during which behavior was continuously recorded. ANY-maze software was utilized to measure locomotor parameters, including total distance traveled, average speed, time spent in the center versus periphery, number of center entries, and immobility duration. These metrics enabled simultaneous evaluation of locomotion and anxiety-related responses, as rodents tend to avoid the center zone in open field tests.

To minimize olfactory confounds, the arena was cleaned with 70% ethanol and air-dried between trials. Randomization of between-group and within-group testing was achieved via pre-assigned animal IDs to prevent systematic bias. Experimenters conducting tests and analyzing data were blinded to treatment group and animal sex.

No animals were excluded from analysis in the OFT, as all completed the full test without technical difficulties or abnormal behavior (e.g., freezing, wall climbing, or jumping). Mice that showed excessive grooming or signs of illness prior to testing would have been excluded, but no such exclusions were necessary for the OFT cohort.

Data were analyzed in a sex-agnostic manner for the primary report, with follow-up sex-stratified comparisons provided in supplemental tables. One-way ANOVA was used to test for group effects, followed by post hoc comparisons when appropriate. The absence of a significant group difference in total distance traveled across treatment conditions confirmed that deficits observed in other behavioral tasks (e.g., EPM or NOR) were unlikely to be secondary to gross motor impairments.

##### 1.3.2. Novel Object Recognition (NOR)

The Novel Object Recognition (NOR) test assessed short-term recognition memory in adolescent mice after perinatal exposure to either single heavy metals (Pb, As, Cd, Cr) or a combination (PACC). The protocol was based on validated methods for juvenile C57BL/6J mice, with adjustments for developmental stage and strain [4-6]. Testing occurred under dim red light during the dark phase, with mice habituated in the testing room for at least 60 minutes beforehand. All equipment was scent-free, and experimenters were blinded to exposure groups throughout the process.

###### Apparatus

The NOR arena was a 40 cm cubic opaque acrylic box with a smooth floor, placed in a sound-dampened chamber under low, even lighting. Video and behavior were recorded with overhead infrared cameras and ANY-maze software. Each trial used two identical and one novel object, pre-validated for equal interest in controls. All objects were non-porous, non-toxic, and 3–4 cm in size to promote exploration without climbing.

###### Procedure

The NOR test consisted of three phases conducted across two consecutive days:

1. **Day 1, Habituation Phase:** Each mouse was individually placed in the empty arena for a 5-minute session to allow unrestricted exploration. No objects were introduced during this period. Mice displaying freezing, excessive grooming, or immobility for more than 50% of the session were identified for further observation but were only excluded if such behaviors impeded standard exploratory activity.

2. **Day 2, Training Phase:** Each mouse was placed in the arena with two identical objects in opposite corners (10 cm from the walls) and allowed to explore for 5 minutes. Object interaction was recorded only when the mouse actively sniffed within 2 cm; climbing or passing by without sniffing was not counted.

3. **Day 2, Testing Phase (30 min post-training):** One of the previously presented objects was substituted with a novel object differing in shape, color, and texture. Mice were permitted to explore the arena for 5 minutes. The arena, floor, and

objects were cleaned with 70% ethanol between trials. Object placement (left/right) was counterbalanced among animals to reduce potential side bias.

**Outcome Measures**

The primary outcome was the **discrimination index**, calculated as:

Discrimination Index (DI):

$$DI = \frac{(Tnovel - Tfamiliar)}{(Tnovel + Tfamiliar)}$$

Where *Tnovel* is the time spent exploring the novel object, and *Tfamiliar* is the time spent exploring the familiar object. Total object exploration time was also recorded. Animals that explored both objects for a combined time <5 seconds were excluded from analysis (<2% of the cohort), consistent with recommendations from prior validation studies.

**Controls and Randomization**

Trials were video recorded and scored by blinded experimenters. Testing order, object identity, and placement were pseudo-randomized. Handling was minimized between phases. Only animals with complete data were included in analyses. Sedatives and food deprivation were not used.

**Interpretation and Relevance**

The NOR task, which leverages rodents' innate preference for novelty without training or rewards, is well-suited for developmental studies. In this study, PACC metal mixture-exposed mice showed significantly reduced novel object preference, indicating impaired short-term recognition memory.

**1.3.3. Elevated Plus Maze (EPM)**

The Elevated Plus Maze (EPM) was used to evaluate anxiety-like behavior in mice after perinatal exposure to specific heavy metals or their mixture (PACC). This test relies on rodents' tendency to avoid open elevated areas, balanced by their exploratory behavior. The protocol was based on established methods validated in juvenile and adult C57BL/6J mice [7,8], and was adjusted for the developmental stage and aims of this study.

All procedures took place during the dark phase of a reversed light cycle (9:00 AM–9:00 PM lights off) under dim red light to minimize stress and replicate nocturnal conditions. Mice were acclimated to the testing room for at least one hour prior to assessment. Experimenters were blinded to the exposure condition, and all testing materials were ensured to be free from residual olfactory cues.

##### Apparatus

The elevated plus maze (EPM) apparatus comprised four arms arranged in a plus configuration, raised 50 cm above the floor. Two opposite arms were enclosed (30 × 5 × 15 cm), while the remaining two arms were open (30 × 5 cm). All arms converged at a central square platform measuring 5 × 5 cm. The maze was constructed from matte black acrylic and mounted on a stable base designed to minimize vibrations. The entire apparatus was situated within a sound-attenuating, low-light chamber and equipped with an overhead infrared camera (Stoelting ANY-maze system) to enable real-time behavioral tracking and analysis.

##### Procedure

Each mouse was placed gently at the center of the maze, facing one of the open arms, and allowed to explore freely for 5 minutes. The arena was cleaned thoroughly with 70% ethanol between animals and allowed to air dry to eliminate scent cues.

The following parameters were recorded automatically via ANY-maze:

- Time spent in open arms (seconds and % of total time)
- Number of entries into open and closed arms
- Total distance traveled and speed
- Time spent in the center zone

Entries were defined as instances when all four paws of the mouse crossed into an arm. Mice that did not leave the center platform within one minute were excluded and given one additional trial. No mice required retesting under this criterion. Animals displaying behaviors such as excessive grooming, freezing for more than 30% of the session, or falling off the maze would have been excluded from the study; however, none of these behaviors occurred.

##### Interpretation and Controls

Increased time or entries into open arms are interpreted as decreased anxiety-like behavior, while a preference for closed arms reflects elevated anxiety. In this study, the PACC metal mixture-exposed mice exhibited significantly reduced open arm time and fewer open arm entries, consistent with an anxiogenic effect of perinatal metal exposure.

To control for confounding motor effects, total locomotor activity (distance traveled, speed) was analyzed. The absence of differences across groups in total movement confirmed that observed anxiety phenotypes were not due to general hypoactivity or sedation.

##### Randomization and Bias Control

Animals were tested in a randomized order within treatment cohorts to prevent circadian or handler bias. The apparatus orientation was rotated every 3–5 animals to mitigate spatial bias due to room geometry or odor trails. All scoring and data interpretation were conducted by personnel blinded to group identity.

##### 1.3.4. Y-maze Spontaneous Alternation

The Y-maze spontaneous alternation test was utilized to evaluate spatial working memory in mice that were perinatally exposed to either individual heavy metals or the PACC metal mixture. This assessment leverages rodents' natural exploratory tendencies, particularly their preference for entering a novel arm over returning to one previously visited, which reflects functional short-term memory. The protocol adhered to established methodologies for juvenile C57BL/6J mice [9,10], with adaptations for minimal handling and streamlined high-throughput video analysis.

All behavioral testing was conducted during the animals' active (dark) phase under dim red lighting, following a minimum one-hour habituation period in the behavior room. At all experimental phases, including testing and analysis, the experimenter remained blinded to treatment group assignments.

##### Apparatus

The Y-maze comprised three arms, each measuring 30 cm in length, 5 cm in width, and enclosed by 15 cm high walls, arranged at 120° angles from a central triangular zone. The apparatus was constructed from matte black acrylic to reduce reflections and glare. The maze was situated within a sound-attenuated, low-light chamber equipped with overhead infrared video monitoring (ANY-maze, Stoelting Co.) for automated behavioral tracking. The central area, where the three arms joined, was digitally delineated in the software to accurately record transitions between arms.

##### Procedure

Each mouse was placed in the center of the maze facing one predetermined arm and allowed to explore freely for 5 minutes. No prior training, rewards, or deprivation protocols were used, and animals were not familiarized with the maze beforehand to preserve the novelty-driven aspect of the test. Between each trial, the maze was thoroughly cleaned with 70% ethanol and allowed to air-dry.

Arm entries were defined as the placement of all four paws into an arm. The sequence of entries was used to calculate the **percent spontaneous alternation** using the standard triad-based formula:

$$Alternation\,\% = \frac{Number\ of\ correct\ alternations}{Total\ number\ of\ possible\ alternations} \times 100$$

Where a correct alternation was defined as entries into three different arms consecutively (e.g., ABC, BCA, or CAB). The total number of possible alternations was calculated as the total number of arm entries minus two.

In addition to alternation percentage, total arm entries, movement distance (cm), time spent in each arm, and time in the center zone were recorded to assess general locomotor activity and rule out hypoactivity as a confound.

**Quality Controls and Exclusion Criteria**

A minimum of eight total arm entries was required for inclusion in the analysis to ensure that spontaneous alternation rates accurately reflected exploratory behavior. Mice with fewer entries were excluded from analysis (representing less than 3% of all animals). No exclusions were made due to technical artifacts or behavioral abnormalities, such as freezing or circling. Testing was conducted under conditions of minimal noise and vibration to eliminate external disruptions.

Scoring procedures were fully automated and validated through manual review of a subset of trials to confirm accuracy. The orientation of the apparatus was systematically alternated across trials to minimize potential spatial cueing or directional bias.

**Interpretation**

Elevated alternation rates are indicative of intact spatial working memory, whereas lower alternation rates suggest potential deficits in short-term spatial encoding or working memory retrieval processes. In the present study, no statistically significant differences in alternation rates were detected among the treatment groups, including the PACC metal mixture group, suggesting that spontaneous alternation behavior remained stable despite the presence of other behavioral and synaptic alterations.

**Methodological Considerations**

Recent work[9] emphasizes the importance of arm visit threshold definitions for accurate alternation scoring. In our implementation, we used the conservative criterion of full-body entry (all four paws) to minimize ambiguity, consistent with standard preclinical practice. The center zone boundary was defined via software coordinates corresponding to the midpoint between arm walls to ensure consistency in transition detection.

##### 1.4. Electrophysiology

###### 1.4.1 Brain Slice Preparation

Electrophysiological recordings were performed on acute coronal brain slices prepared within 72 hours after behavioral testing. Mice were anesthetized with isoflurane, decapitated, and their brains quickly extracted and placed in cold, oxygenated sucrose-based cutting solution. Coronal slices (300 µm) from the mPFC (+1.8 to +1.6 mm from bregma) and dorsal CA1 (−1.5 to −2.0 mm from bregma) were cut using a VT1200S vibratome.

Slices recovered in ACSF at 32–34°C for 45 minutes and were then kept at room temperature until recording. ACSF was continuously bubbled with 95% $O_2$/5% $CO_2$. Table S2 lists the salts and other chemical compounds that constitute the different solutions utilized during the recording.

Tissue health was prioritized during slicing and incubation. Slices were oriented along the somatodendritic axis based on anatomical landmarks, and all dissection tools were precooled. Unlike some protocols, transcardial perfusion was not used. Recovery time was optimized to maintain cell integrity without inducing artificial plasticity.

###### 1.4.2 Recording Procedure

Recordings were performed in a continuously perfused submersion-type chamber containing ACSF maintained at 32°C (flow rate: 1–2 mL/min), with temperature regulation provided by a TC-344C inline temperature controller (Warner Instruments). Neurons were visualized using infrared differential interference contrast (IR-DIC) optics with a 40× water-immersion objective on an Olympus BX51WI microscope. Recordings focused on layer II/III pyramidal neurons within the mPFC and CA1 pyramidal neurons.

Borosilicate glass pipettes (resistance 4–6 MΩ) were pulled with a Sutter P-1000 puller and filled with a potassium gluconate-based internal solution. Patch pipettes were maintained under positive pressure to prevent tip obstruction. Upon contact with the cell membrane, slight suction was applied to achieve gigaseal formation, followed by brief pulses of negative pressure to rupture the membrane and establish whole-cell configuration. Series resistance and input resistance were monitored throughout each recording; cells exhibiting a change in series resistance greater than 20% were excluded from further analysis.

###### 1.4.3 Electrophysiological Measurements

**Intrinsic Excitability**

Whole-cell current-clamp recordings were used to assess neuronal excitability. Square-step current injections with 100 ms duration, ranging from −50 to +200 pA in 25 pA increments, were applied from a holding potential of −60 mV. The number of action potentials generated per step was recorded and utilized to construct excitability profiles. Variations in spike number per step indicated modifications in intrinsic excitability. Series resistance compensation was not used; all cells were monitored to ensure passive membrane stability.

This approach enabled comparison of the intrinsic firing properties of pyramidal neurons between groups and assessment of cell-intrinsic changes in excitability that may be related to behavioral phenotypes associated with metal exposure. The chosen current step range was based on prior studies in cortical neurons and followed guidelines from established patch-clamp electrophysiology protocols.

**sEPSCs**

Spontaneous excitatory postsynaptic currents were evaluated by conducting recordings in voltage-clamp mode, with the membrane potential maintained at −70 mV. Under these conditions, AMPA receptor-mediated sEPSCs were measured without the use of pharmacological blockers. Event detection and subsequent analysis were performed offline using Clampfit 11.2 (Molecular Devices), employing a threshold determined by the baseline noise level. Parameters such as amplitude, frequency, and decay kinetics were quantified, and all detected events were manually verified to ensure data integrity and to exclude artifacts or overlapping signals. The sEPSC analysis enabled an assessment of synaptic-level modifications in glutamatergic transmission resulting from perinatal exposure to metal mixtures.

**Table S2. Details of the electrophysiological solutions.**

| **Component** (mM) | **Cutting Solution** (Protective) *+ | **Standard ACSF** (Recovery and Recording) *+ | **Internal Solution** (Pipettes for whole-cell recordings) *+ |
|---|---|---|---|
| **NaCl** | — | 125 | 2.5 |
| **Sucrose** | 230 | — | — |
| **KCl** | 2.5 | 2.5 | 2.5 |
| **$NaHCO_3$** | 25 | 25 | — |
| **$NaH_2PO^4.H_2O$** | 1.25 | 1.25 | — |
| **Glucose** | 25 | 25 | — |
| **$MgCl_2.6H_2O$** | 1 | 1 | 2 |
| **$CaCl_2.2H_2O$** | 1 | 1 | — |
| **HEPES** | — | — | 10 |
| **K-Gluconate** | — | — | 145 |
| **Mg-ATP** | — | — | 2 |
| **Na-GTP** | — | — | 0.5 |

* **pH:** For Cutting solution and standard ACSF: 7.2–7.4, for internal solution: 7.2–7.3 (all adjusted with KOH)

+ **Osmolarity (mOsm):** For Cutting solution and standard ACSF: 300–310, for internal solution: ~285

**2. Supplemental Figures (S1-4) and Supplemental Tables (S3 and S4)**

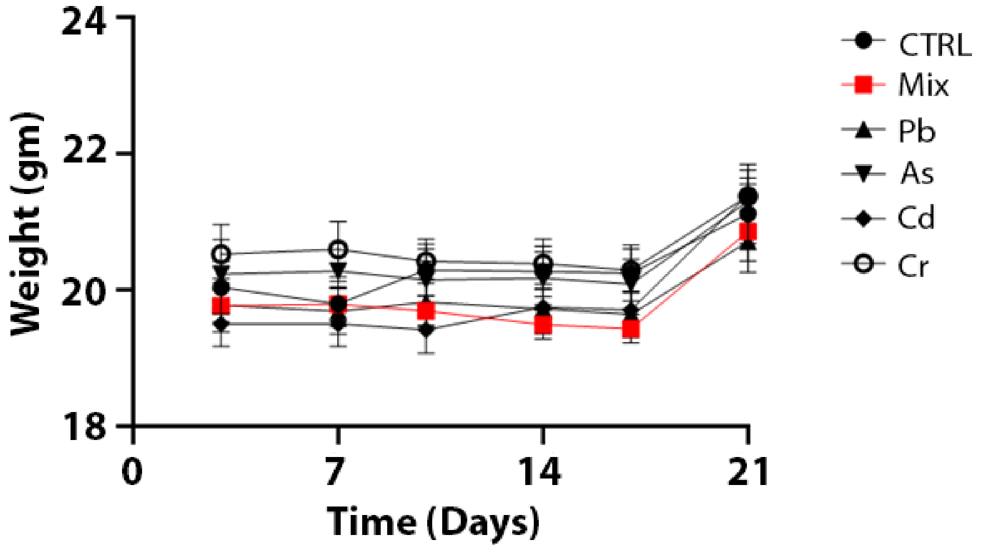


**Figure S1. Perinatal exposure to PACC metal mixture or individual metals does not significantly affect maternal body weight during gestation and lactation.** Maternal body weight was monitored over the course of exposure to either control water (CTRL), PACC metal mixture (Mix; Pb, As, Cd, Cr(VI)), or individual metals. No significant differences in body weight were observed among the groups at any time point from preconception through weaning (Days 0–21). Data are presented as mean ± SEM.

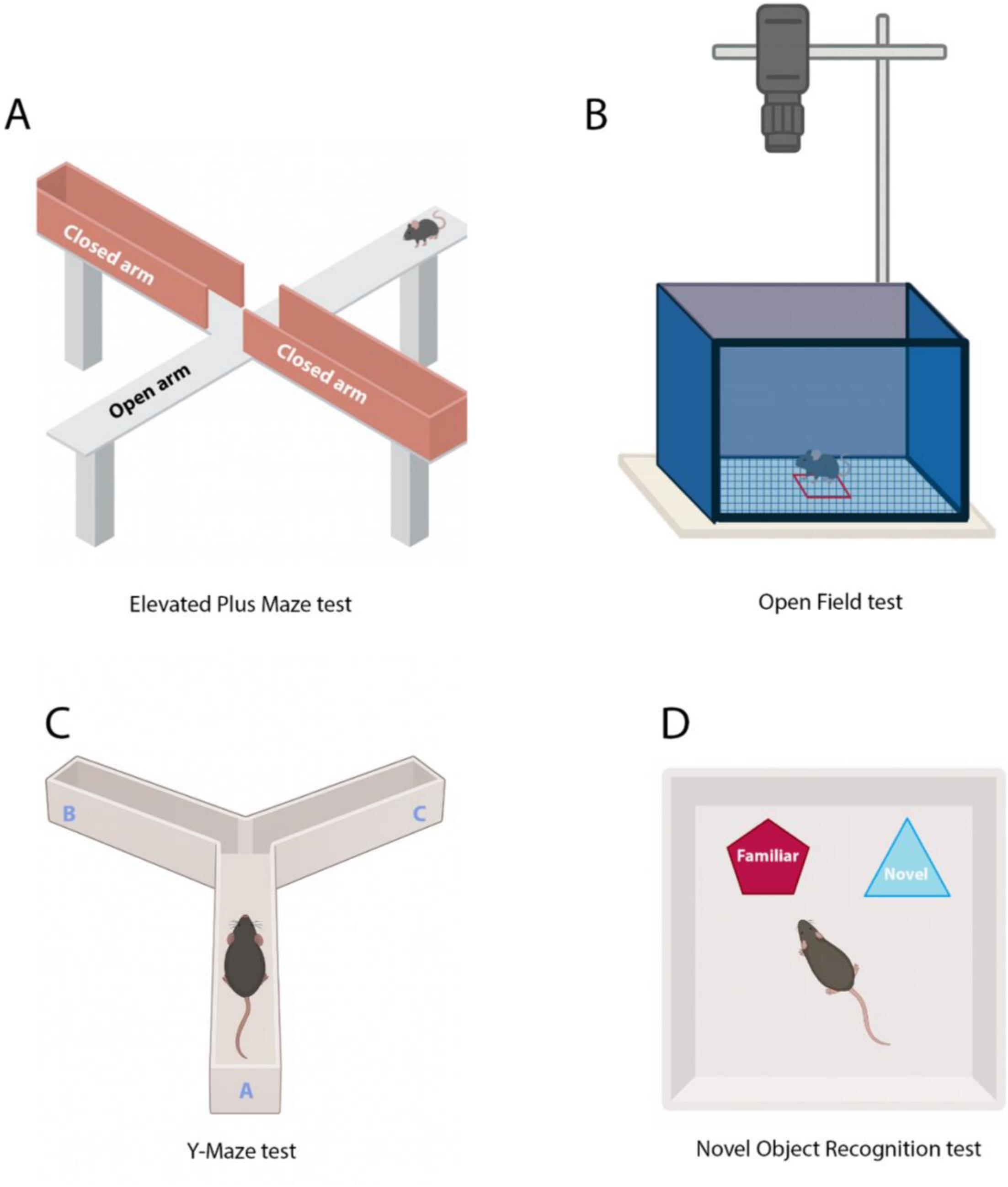


**Figure S2. Schematic representation of behavioral paradigms used to assess neurocognitive outcomes following perinatal metal exposure.**

**(A)** Elevated Plus Maze (EPM) test was used to evaluate anxiety-like behavior by measuring time spent in open vs. closed arms. **(B)** Open Field Test (OFT) assessed locomotor activity and general exploration based on total distance traveled in an open arena. **(C)** Y-Maze test was employed to measure short-term spatial working memory by quantifying spontaneous alternation behavior. **(D)** Novel Object Recognition (NOR) test evaluated short-term recognition memory by comparing exploration time of novel vs. familiar objects. These paradigms were conducted under dim red light during the active (dark) cycle, and outcomes were used to determine the impact of perinatal exposure to PACC metal mixture (Pb, As, Cd, Cr(VI)) metal mixtures on behavioral function in juvenile mice.

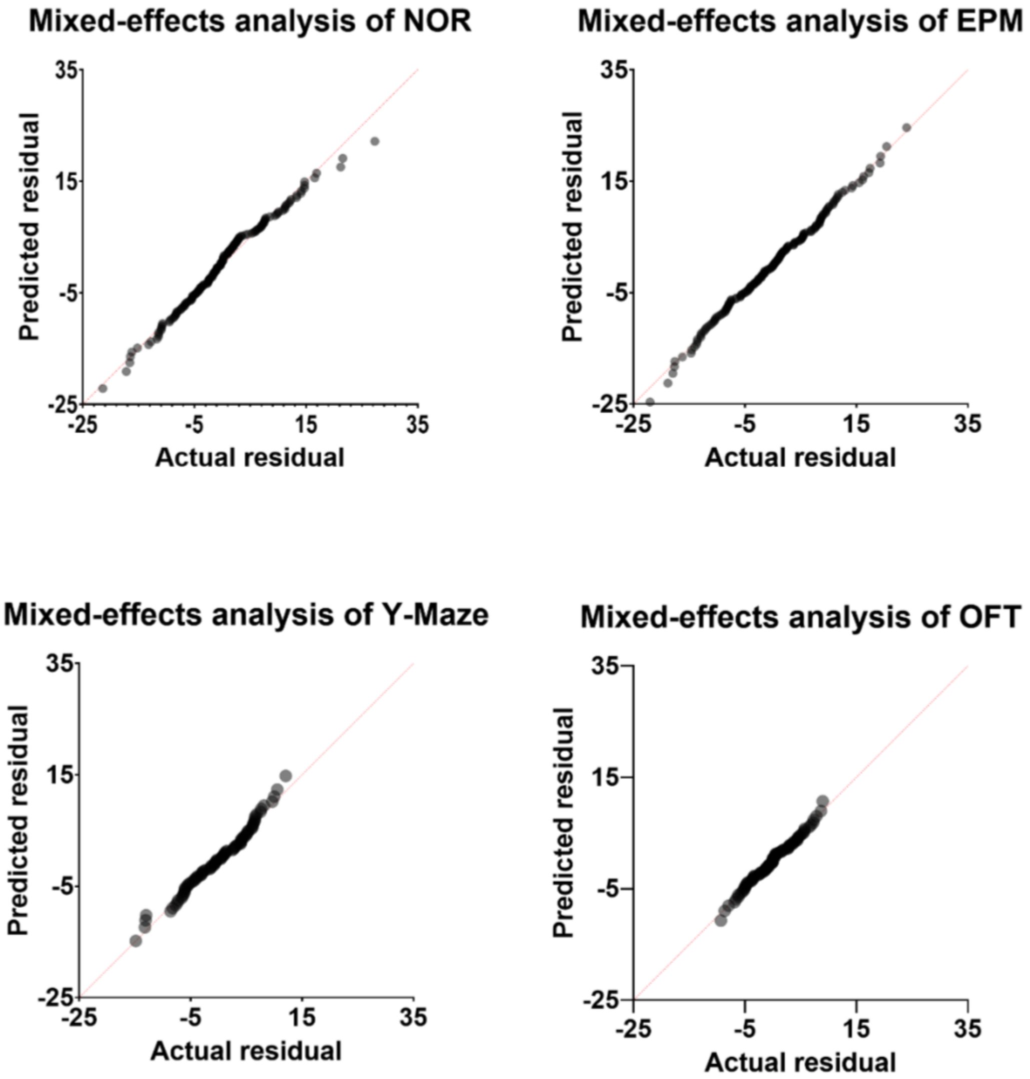


**Figure S3. Residual QQ plots for mixed-effects models applied to behavioral test data.**

Quantile-quantile (QQ) plots show the distribution of residuals from mixed-effects models used to analyze behavioral outcomes in: (top left) Novel Object Recognition (NOR), (top right) Elevated Plus Maze (EPM), (bottom left) Y-Maze, and (bottom right) Open Field Test (OFT). In each case, the residuals follow a near-linear trend, suggesting that model assumptions (e.g., normality of residuals) were adequately met and validating the use of mixed-effects analysis for these behavioral paradigms.

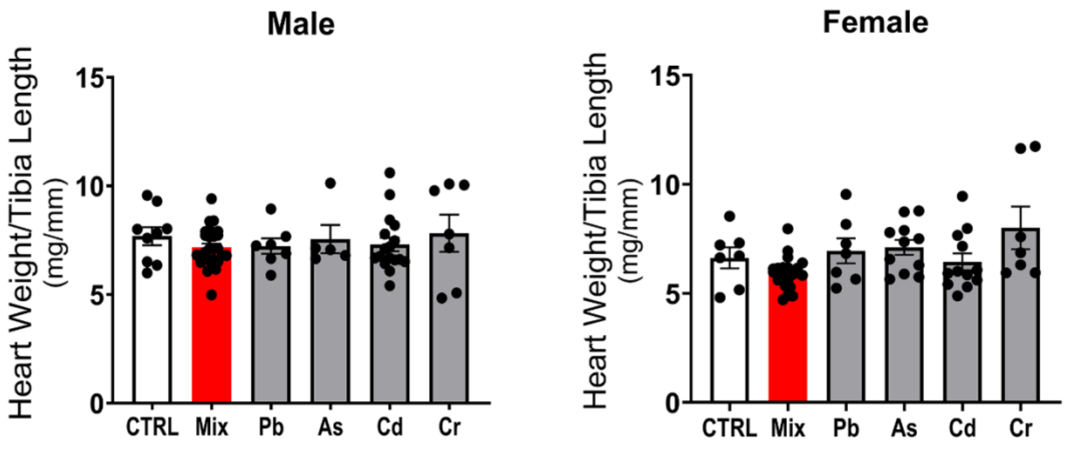


**Figure S4. Perinatal exposure to individual metals or PACC metal mixture does not significantly alter normalized heart size in offspring.**

Heart weight normalized to tibia length (mg/mm) was measured in juvenile male (left) and female (right) C57BL/6J mice perinatally exposed to control water (CTRL), PACC metal mixture (Mix; Pb, As, Cd, Cr(VI)), or individual metals. No significant differences were observed between exposed groups and controls in either sex. Although male offspring exposed to cadmium (Cd) showed a trend toward reduced heart size, this effect coincided with decreased body weight and did not reach statistical significance. Error bars represent mean ± SEM.

**Table S3. Perinatal exposure to the individual metal and metal mixture (PACC) shows no change in the Open field and Y-maze alternation, Results are mean ± SEM**

| | Exposure | | | | | |
|---|---|---|---|---|---|---|
| Test | Unexposed | PACC metal mixture | Pb | As | Cd | Cr |
| OFT | 18.78±1.253 | 20.9±1.258 | 15.82±1.767 | 19.47±1.016 | 21.55±0.4868 | 17.8±0.9172 |
| Y-Maze | 57.54±2.493 | 52.65±1.159 | 53.07±2.099 | 53.13±1.481 | 57.74±1.939 | 54.7±1.502 |

**Table S4. Results of NOR and EPM, separated by males and females, results are mean ± SEM**

| | **Males** | | | | | |
|---|---|---|---|---|---|---|
| Test | Exposure | | | | | |
| | Unexposed | PACC metal mixture | Pb | As | Cd | Cr |
| NOR | 62.83±1.657 | 53.8±1.332 | 56±3.612 | 57.94±4.189 | 57.22±1.517 | 60.49±2.32 |
| EPM | 38.01±2.881 | 27.03±1.562 | 20.77±2.164 | 29.94±3.653 | 30.98±2.075 | 37.05±3.948 |
| | **Females** | | | | | |
| | Exposure | | | | | |
| | Unexposed | PACC metal mixture | Pb | As | Cd | Cr |
| NOR | 64.21±2.401 | 51.81±1.796 | 58.8±3.531 | 55.13±2.289 | 57.77±1.739 | 54.64±1.644 |
| EPM | 38.01±2.881 | 27.03±1.562 | 20.77±2.164 | 29.94±3.653 | 30.98±2.075 | 37.05±3.948 |

#### 3. Method for Abbreviated Additivity Analysis

##### 3.1. Background

This study investigates individual and mixture exposures to PACC metals and effects on neurological outcomes. Doses for PACC metals were selected from National Primary Drinking Water Regulations which are legally enforceable standards developed by EPA to control contaminants in drinking water. The standards selected are maximum contaminant levels (MCLs) for Ar, Cr, Cd and are derived from existing reference doses (RfDs) [11]. EPA's RfDs are estimates of daily exposure that are likely to be without appreciable risk of harmful effects over a lifetime, including for sensitive populations [12] Pb is a known developmental neurotoxicant and does not have an RfD as there is no known threshold of exposure that is considered to be safe [13]. The Pb standard is therefore not an MCL, it is an action level (AL) that if exceeded requires the water system to take actions such as service line replacement [14]. Health outcomes of concern associated with the RfDs for Ar, Cr, and Cd include adverse effects on the kidneys, skin, and circulatory system. However, evidence indicates that Ar, Cr, Cd, and Pb all share a common mode of action (MOA) for neurological impairment; PACC can generate reactive oxygen species or impair calcium homeostasis, leading to inhibition of normal developmental signaling pathways, altered synaptic plasticity, glial activation, and impaired mitochondrial function [15-19].

EPA uses two basic approaches for predicting the effects of a mixture, dose addition and response addition [20]. When mixture effects are additive, there are no interactions among chemicals in the mixture. If there are interactions, effects could be synergistic (i.e., greater than additive) or antagonistic (i.e., less than additive). The use of dose addition to predict mixture effects relies on three main assumptions [20]. First, the chemicals in the mixture are toxicologically similar and act through the same mode of action or on the same target organs. Second, since the chemicals are toxicologically similar, dose-response curves are parallel, and mixture components can be considered dilutions of each other. Third, the mixture response can be predicted by the sum of the scaled doses for individual chemical components. If the chemicals in the mixture act through different MOAs, or "independent action", the response to a single chemical in the mixture is not influenced by the presence of a second chemical. If independent action is assumed for the chemicals in the mixture, EPA may use response addition [20]. Methods include summing risk estimates for the individual chemicals if effects are statistically independent, or effect summation which sums the measured responses of the mixture components. No assumptions are made about the shapes of dose-response curves for component chemicals for response addition.

We use these two basic approaches, dose addition and response addition, to perform a simplistic additivity assessment. To assess additivity using dose addition, we adopted a four-step approach for evaluating dose additivity in mixtures proposed by Hertzberg et al. [21]. This approach uses relative potency factors (RPFs), which express the potency of each chemical relative to a chosen "index chemical". This method is widely used in regulatory contexts and is well-suited when chemicals act via the same MOA and full dose-response data are available. The MOA shared by the PACC metals satisfies a key assumption needed to assess dose additivity, that the metals are toxicologically similar. Full dose-response data are not available, and we are limited to effects observed from PACC metals at single doses. This is a major limitation as applied to this study. However, to enable a simple additivity analysis, we assume similar dose-response curves for the PACC metals and that the effects levels observed at the applied doses are equivalent to lowest observed effect levels (LOELs). (RPFs have been calculated with LOELs, NOELs, $LD_{50}$s, and other benchmarks [22]. We also applied a simple method for assessing response additivity [20,23]. This approach assumes independent action of the chemical, an assumption that may not hold considering the shared MOA of the PACC metals. Future studies supplying additional dose-response data can update the rudimentary estimates provided here.

##### 3.2. Methods

Dose addition and response addition approaches to assessing additivity were only applied to behavioral tests with significant results, Elevated Plus Maze (EPM; anxiety-like behavior) and Novel Object Recognition (NOR; short-term memory), for all mice combined (male and female mice). Each metal was tested individually at a single dose level (the MCL or AL), and as a mixture with each metal at the same concentration (refer to main text for a full description of methods).

###### 3.2.1. Dose additivity

A simplified dose addition approach [21] was applied to evaluate the mixture effect of the PACC metal mixture. The metals in the PACC metal mixture are assumed to be toxicologically similar because they share an MOA that may lead to adverse neurological effects [15-19]. Assuming toxicological similarity, the metals can be considered dilutions of each other and the relative potency factor approach can be applied (with the limitations previously noted).

First, net responses are calculated for behavioral performance tests, Elevated Plus Maze (EPM; anxiety-like behavior) and Novel Object Recognition (NOR; short-term memory), by subtracting the observed response from the control response. Second, the mixture concentration is expressed in terms of the index chemical. Pb is selected as the index chemical because it has the strongest body of evidence supporting neurodevelopmental effects, and relative potency factors (RPF) were calculated for As, Cd, and Cr as the ratio of the index chemical dose to the dose of the other metals:

$$RPF_i = \frac{D_{index}}{D_i}$$

$RPF_i$ = relative potency factor
$D_i$ = dose (mg/kg/day) of metal i
$D_{index}$ = dose (mg/kg/day) of Pb, the index chemical (RPF = 1)

The dose of As, Cd, and Cr were multiplied by their RPFs to convert into an equivalent dose of the index chemical. The total index chemical equivalent dose (ICED) of the PACC metal mixture was then calculated as the sum of the index-equivalent doses:

$$ICED_{mixture} = \sum_{i=1}^{n} D_i * \ RPF_i$$

Third, the mixture response is predicted. To predict the mixture response, a scaling factor *c* was first estimated using the net response to the index chemical Pb and the dose:

$$c = \frac{Net\ Response_{index}}{D_{index}}$$

Then, the constant was used to predict the net response of the PACC metal mixture:

$$Predicted\ Net\ Response = c * ICED_{mixture}$$

Fourth, the predicted mixture response from the third step is compared to the observed mixture response. A positive difference may indicate synergism, a negative difference may indicate antagonism, while no difference may indicate the mixture effects are additive.

##### 3.2.2 Response additivity

The response addition approach makes no assumptions about the shapes of the dose response curves for the metals in the PACC metal mixture, but also assumes "independent action" or that the metals act on the neurological endpoint through different, independent modes of action[20]. This assumption may not hold, given the evidence that PACC metals are toxicologically similar, and is a major limitation of this approach. The response addition method, where risk estimates for each chemical component in the mixture are summed to predict the mixture effect, was originally developed for mortality or binary response data and is most commonly applied in cancer risk assessments [24,25]. The method has been adapted for continuous data scaled to the range [0,1] and is used here as a second approach to assessing additivity in the PACC metal mixture [20,23,26].

First, the net response is calculated for the behavioral performance tests, EPM and NOR, by subtracting the observed response from the control response. The expected effect of the mixture is calculated as:

$$E_{mixture} = 1 - \prod_{i=1}^{n}[1 - E(D_i)]$$

Where $E_{mixture}$ is the predicted effect of the mixture and $E(D_i)$ is the observed effect of the individual chemical at dose $D_i$.

##### 3.3. Results

Mean responses from behavioral performance tests with significant results and net responses for male and female mice together are presented in **Table S5**.

**Table S5. Mean (range) behavioral performance test responses for Elevated Plus Maze (EPM; anxiety-like behavior), Novel Object Recognition (NOR; short-term memory), for male and female mice, and net responses (control-observed).**

| **Behavioral Test*** | **NOR** | | **EPM** | |
|---|---|---|---|---|
| **Treatment** | **Mean (range) novel object preference (%)** | **Net response** | **Mean (range) open arm time (%)** | **Net response** |
| Control | 64.49 (45.93–83.05) | - | 34.71 (10.83–58.59) | - |
| Mixture | 52.92 (36.95–68.89) | 11.57 (8.98–14.16) | 25.45 (7.32–43.58) | 9.26 (3.51–15.01) |
| As | 56.31 (37.48–75.14) | 8.18 (8.45–7.91) | 34.04 (10.62–57.46) | 0.67 (0.21–1.13) |
| Cd | 57.46 (44.32–70.60) | 7.03 (1.61–12.45) | 28.2 (10.27–46.13) | 6.51 (0.56–12.46) |
| Cr | 57.56 (43.23–71.89) | 6.93 (2.70–11.16) | 37.77 (12.39–63.15) | -3.06 (-1.56– -4.56) |
| Pb | 57.6 (39.06–76.14) | 6.89 (6.87–6.91) | 25.4 (10.64–40.16) | 9.31 (0.19–18.43) |

*Additivity assessment was only performed for behavioral tests with significant results.

###### 3.3.1. Dose addition (RPF) approach

**Table S6** provides the concentrations of PACC metals administered through drinking water (parts per billion (ppb)), the dose of each metal (mg/kg/day) based on amount of water consumed and bodyweights of mice. The relative potency factors (RPFs) of the metals with Pb as the index chemical are also presented, assuming PACC metals are toxicologically similar.

**Table S6. Experimental exposure concentrations, calculated doses based on water consumption and bodyweights, and relative potency factors (RPFs) of Pb, As, Cd, and Cr.**

| Metal | ppb | Mean (range) dose (mg/kg/day) | • RPF |
|---|---|---|---|
| Pb | 15 | 0.0031 (0.0023–0.0038) | 1 (index) |
| As | 10 | 0.0022 (0.0019–0.0025) | 1.5 |
| Cd | 5 | 0.001 (0.0007–0.0013) | 3 |
| Cr | 100 | 0.021 (0.015–0.028) | 0.15 |

The index chemical equivalent dose (ICED or the lead equivalent dose) of the PACC metal mixture is 0.0125 mg/kg/day. The predicted net mixture responses (**Table S7**) were larger than observed responses for NOR and EPM behavioral tests. The difference indicates a less than additive response or the possibility of antagonism.

**Table S7. Net observed mean PACC metal mixture responses from NOR and EPM tests, predicted PACC metal mixture responses using dose addition and RPFs, and differences between observed and predicted responses.**

| Response | NOR* (%) net mean mixture response | EPM* (%) net mean mixture response |
|---|---|---|
| Observed | 11.57 | 9.26 |
| Predicted | 28.11 | 37.99 |
| Difference | -16.54 | -28.73 |

* Elevated Plus Maze (EPM; anxiety-like behavior), Novel Object Recognition (NOR; short-term memory)

##### 3.3.2. Response addition approach

The predicted net mixture responses (**Table S8**) were also larger than observed responses for NOR and EPM behavioral tests using response addition (assuming independent action). The difference indicates a less than additive response or the possibility of antagonism.

**Table S8.** **Net observed mean PACC metal mixture responses from NOR and EPM tests, predicted PACC metal mixture responses using effect summation, and differences between observed and predicted responses.**

| Response | NOR* (%) net mean (range) mixture response | EPM* (%) net mean (range) mixture response |
|---|---|---|
| Observed | 11.57 (8.98 – 14.16) | 9.26 (3.51 – 15.01) |
| Predicted | 26.02 (18.38 – 33.32) | 13.20 (-0.58 – 26.18) |
| Difference | -14.45 (-9.4 – -19.16) | -3.94 (4.09 – -11.17) |

* Elevated Plus Maze (EPM; anxiety-like behavior), Novel Object Recognition (NOR; short-term memory)

##### 3.4. Conclusions

Results from the simplified dose additivity and response additivity analyses suggest less that additive mixture effects or antagonism. These initial results should be interpreted with caution due to limitations associated with using response data from PACC metals administered in water at a single concentration, assuming that dose-response curves are parallel in the absence of response data at multiple doses for dose additivity analysis. Likewise, assumptions made for response addition analysis, that the PACC metals act independently, may not hold due to similarities in the PACC metals modes of action, and is a major limitation of this approach. These results should be updated when more complete dose-response data becomes available for the PACC metal mixture.
